# Scaling between DNA and cell size governs bacterial growth homeostasis and resource allocation

**DOI:** 10.1101/2021.11.12.468234

**Authors:** Boyan Li, Songyuan Zhang, Le Zhang, Xiaoying Qiao, Yiqiang Shi, Cheng Li, Qi Ouyang, Ping Wei, Long Qian

## Abstract

Bacteria maintain a stable cell size and a certain DNA content through proliferation as described by classic growth laws. How cells behave when this inherent scaling is broken, however, has rarely been interrogated. Here we engineered *Escherichia coli* cells with extremely low DNA contents using a tunable synthetic tool *CRISPRori* that temporarily inhibited chromosome replication initiation. A detailed mechanistic model coupling DNA replication, cell growth, and division revealed a fundamental DNA-centric growth law, which was validated by two observations. First, lineage dynamics were robust to large *CRISPRori* perturbations with division cycles rapidly restoring through a timer mechanism rather than the adder rule. Second, cellular growth transitioned into a linear regime at low DNA-cytoplasm ratios. Experiments and theory showed that in this regime, cellular resource was redirected to plasmid-borne gene expression. Together with the ability of *CRISPRori* to bi-directionally modulate plasmid copy numbers, these findings suggest a novel strategy for bio-production enhancement.

## Introduction

Cell growth is intertwined with the cell’s hereditary stability and size stability. That is, growth has to be constantly in dynamic coordination with the replication of genetic materials and cell division. Eukaryotic cells invest heavily in fail-safe mechanisms to maintain normal growth states. For example, classical DNA replication licensing and completion checkpoint mechanisms throughout the cell cycle ensure the correct chromosomal ploidy in daughter cells after division^1,2^. Likewise, observations have been made for important intracellular pathways that scale or not scale with cell size, with their functional implications conjectured^3–5^. In yeast and mammalian systems, deregulated cell size has been shown to be related to senescence and various pathological situations^5–8^.

Devoid of eukaryotic cell cycle checkpoint and size control mechanisms, the prokaryotic cell serves as a *minimal model* for cell growth regulation. Shifts in growth states underlie bacterial adaptation to the astonishing environmental diversity^9–13^. On the other hand, understanding growth as general cellular physiology has paramount importance for microbial engineering of bio-production and other synthetic biological applications^14^. As cell growth is a direct consequence of the expression of structural proteins, its rate has been shown to have a linear relationship to the ribosomal content^11^, which in turn acts on the general translational level. Therefore, the autocatalytic production of ribosomes is believed to be the major cause of exponential cell growth^15^, with its rate defined by the translation elongation rate. During growth, genome replication and cell division are timely triggered by related proteins^16,17^ to result in macroscale observables such as cell size, DNA content and the membrane surface area.

It is noteworthy that genomic DNA plays multiple roles in these processes. On the one hand, DNA provides the ultimate template for the making of the cellular proteome that collectively drives mass accumulation and volumetric expansion. In this sense, DNA, as well as transcription and translational machineries, which are themselves encoded on DNA, serve as primary catalysts for the cell as a self-replicator. Therefore, gene dosage^18–20^ and the availability of transcription and translational proteins^21–23^ strongly influence the rate of the self-replicating reaction. On the other hand, DNA directly participates in genome replication and cell division. Emerging work on the mechanistic origin of DNA replication initiation has served to validate an ATPase, DnaA, as an initiator protein, whose accumulation at bacterial DNA replication origin *oriC* catalyzes DNA unwinding^16,24,25^. As for cell division, a tubulin homolog FtsZ was identified to polymerize into a Z-ring structure that guides the bacterial cytokinetic apparatus^26,27^. DNA is involved here through a mechanism known as nucleoid occlusion^17^ by providing cues for timing and positioning of Z-rings. Given the broad range of cell size in the prokaryotic kingdom^28^, it is conceivable that the scaling between DNA and growth-related proteins enters the exquisite regulation of cellular growth.

Early observations of bacterial growth yielded phenomenological “growth laws” including the SMK model^10^, the Cooper-Helmstetter model^29^, Donachie’s hypothesis^30^, and the adder law^24,28,31–33^. Modern experimental investigations, especially transcriptomic and proteomic ones, have been conducted mainly under various nutritional or stress conditions^11–13,34–36^. Intracellular resource allocation has been postulated as the central notion for growth regulation and shown to be implemented through global translational regulation as well as transcriptional modulation via the stringent response^37^. Such models were able to explain growth rate changes upon environmental shifts. However, in most growth experiments, cells were not driven into extreme states with their genomic content significantly deviated from the normal range. Thereby, although genome replication was prominently present in the classic growth laws^10,29,30^, DNA largely remained a “hidden” variable in modern mechanistic models. Meanwhile, gene dosage has been discussed in the context of relative gene expression but is hardly connected to growth^18–20,38,39^. Several recent theoretical studies on balanced cell growth have proposed coarse-grained partition models of RNA polymerase or ribosomes to nucleotides to explain changes in growth rates^23,40^, but experimental verification in prokaryotes has not been reported. It remains an interesting question as to how much one could perturb cellular physiology in terms of DNA content and its scaling with size and other proteins, and under these situations, whether growth laws hold true or require modification.

In this work, we constructed a synthetic dCas9 based system, *CRISPRori*, to inhibit genomic replication initiation in *E. coli* cells. The system was able to reduce the DNA-cytoplasm ratio in *E. coli* cells to an extent far below the normal range. With this experimental system, we showed that homeostasis of cellular growth is maintained through lineage dynamics even when cells were driven away from their native state. A multiscale model was constructed to interrogate the molecular interactions involved in coordinated genome replication, cellular growth and cell division. The model and experimental results revealed the central role of DNA content in cellular resource allocation and the maintenance of exponential growth.

## Results

### A CRISPR-based system decoupled chromosome replication from cell growth

To directly perturb DNA replication, we constructed a synthetic replication initiation control system, *CRISPRori*, in which dCas9 was programmed to target the *oriC* region of *E. coli* (**Fig. 1A**). The system was integrated into a single medium-copy-number plasmid, with dCas9 driven by the arabinose-inducible promoter pBAD and the single guide RNA (sgRNA) driven by the constitutive promoter J23119 **(Fig. 1B)**. The inhibitory effect of dCas9 was tested using the CRISPR interference experiment, where the expression of target mRFP was dampened over 30-fold compared to the negative control with non-complementary sgRNA (a 20-nt poly-adenine sequence), under the saturating concentration of arabinose. We found the attachment of an ssrA degradation tag to the C-terminus of dCas9 was helpful in obtaining a moderate inhibitory strength, enabling a larger dynamic range of suppression with respect to inducer concentrations **(Supplementary Fig. 1)**. Therefore, the dCas9-ssrA fusion gene was used in the following experiments.

**Figure 1.**
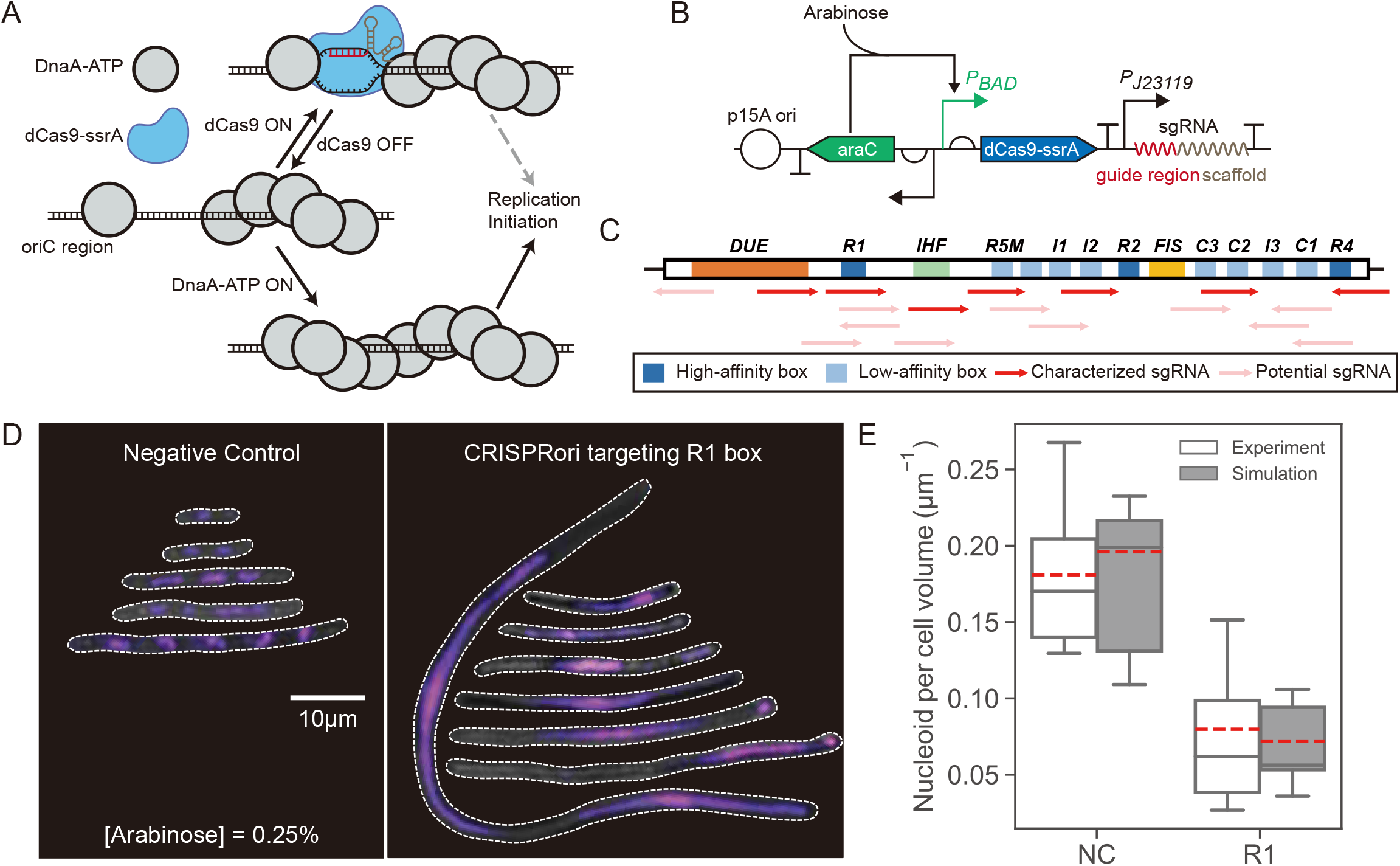
Design of *CRISPRori* to perturb prokaryotic genome replication initiation. **A.** A schematic diagram of the synthetic replication initiation perturbation system, *CRISPRori*. dCas9 is programmed to target various binding sites of DnaA-ATP in the *oriC* region. **B.** Plasmid construction of *CRISPRori*. dCas9-ssrA and sgRNA is integrated on a single plasmid. Expression of dCas9-ssrA is induced by arabinose, while the sgRNA is transcribed constitutively. **C.** An sgRNA library (red and pink arrows) targeting different sites on *oriC*. While many targetable sequences exist in the *oriC* region, seven were experimentally characterized (red arrows). Characterized sgRNAs cover all categories of DNA elements, including (i) DnaA-boxes with high (dark blue) and low (light blue) affinities to DnaA-ATP, respectively; (ii) DNA unwinding element (DUE, orange box) where replisomes bind; and (iii) binding site of the regulatory protein IHF (green box). **D.** *CRISPRori* decoupled DNA replication from cell growth. Confocal images of cells expressing *CRISPRori* with sgRNA targeting the R1 box or a negative control sgRNA (20-nt poly(A)). Nucleoids were stained by DAPI (purple). The inducer concentration was 0.25%. **E.** Experimental and simulation results of number of nucleoids per unit cell length. In the NC group, cells occasionally grew very long but maintained a certain nucleoid-to-cytoplasmic ratio (~0.1592 μm^−1^). In the *CRISPRori* group, cells grew longer without replicating their chromosomes, resulting in much lower nucleoid-to-cytoplasmic ratios (0.03196 μm^−1^). *N*_cell_ = 30~50. In this and all box plots below, the gray lines in the middle of boxes show the median, the red dashed lines show the mean, the boxes show the upper and lower quartiles and the whiskers extend through 10%-90% of the data.

The *oriC* region of the *E. coli* genome carries multiple specific binding sites for DnaA and other accessory proteins for replication initiation^16,25,41^. To test *CRISPRori*’s ability to suppress replication initiation, an sgRNA library collectively covering nearly 90% of the *oriC* region was constructed **(Fig. 1C, Supplementary Table 1)**. These sgRNAs were pre-screened *in silico* to minimize off-target effects^42^.

Induction of the *CRISPRori* system led to up to 25-fold elongated cell morphologies **(Fig. 1D)**. Elongation occasionally occurs in normally growing *E. coli*^43^ or *E. coli* under certain stress conditions^44^. However, unlike under the latter conditions where cells exhibit multi-nucleoid “bean-like” structures without significant changes in the DNA-cytoplasm ratio^45^, *CRISPRori* perturbed cells retained a single nucleoid region as shown by fluorescent staining, suggesting that genome replication had been effectively decoupled from cellular growth **(Fig. 1D)**. DNA-cytoplasm ratios, as measured by the number of distinct nucleoid regions divided by cell lengths, were 0.1810 μm^−1^ and 0.0798μm^−1^ for the negative control and *CRISPRori* inhibited cells, respectively **(Fig. 1E)**. Secondly, the elongated cell morphologies were not associated with an irreversible transition into a reduced growth state such as senescence. Cells were able to restore their normal morphology between episodes of *CRISPRori* induction **(Supplementary Fig. 2D)**.

### Division size and adder were sensitive to three tunable ways of initiation perturbation

To monitor cell morphology under controllable environments, we used time-lapse microscopy and a well-designed microfluidic chip^46^ **(Supplementary Fig. 2A,B)** with chambers of 90 μm in width and 1.2 μm in height where *E. coli* cells freely grew in a single layer. A set of image analysis methods enabled us to acquire quantitative cell growth data for 10 hours, with high spatial (0.1 μm) and temporal (3 min per frame) resolutions, on both single-cell and population levels (**Methods**).

As in the pilot test, cellular morphological changes were most conspicuous due to *CRISPRori* induction **(Fig. 1D, Supplementary Fig. 2C)**. For rod-shaped bacteria, cell size can be estimated by cell width and length. Both data were extracted from time series of microscopic images of *CRISPRori* perturbed *E. coli* populations. Under all conditions, we found minimal or no significant change in cell widths compared to the control group **(Supplementary Fig. 3)**. Cell width has been previously associated with chromosome elongation and increased nucleoid complexity during bacterial genome replication^47^. This suggested that *CRISPRori* indeed held replication at the initiation stage. Subsequently, cell length was used as a proxy for cell size.

Cell lengths right before division (i.e., division lengths) changed with varying strengths of *CRISPRori* activities, which we were able to modulate in three ways: (1) by targeting *oriC* boxes with differential affinities to the major replication initiation protein DnaA, (2) by changing the inducer concentration and (3) by changing the sgRNA length.

There are eleven sites in the *oriC* region on the *E. coli* chromosome for the specific binding of DnaA, of which three (R1, R2, and R4) are high-affinity binding sites, and the rest (R5M, τ2, I1, I2, C1, I3, C2, and C3) are low-affinity ones^16,25,41^ **(Fig. 1C)**. Our results showed that the strong-to-weak ordering of *CRISPRori* inhibitory strengths was R1, R5M, R4, C2, and I2 (the rest were not covered by the sgRNA library), with the average division lengths ranging from less than 5 μm to over 20 μm **(Fig. 2B, Supplementary Fig. 4A)**. The notable variations among target boxes suggested *CRISPRori* did not work as a roadblock to the replication fork, consistent with previously observed molecular kinetics of dCas9 on the *E. coli* genome^48^. Rather, it worked presumably by interfering with pre-initiation processes. To model replication initiation under *CRISPRori* inhibition, we noted two observations. First, although the dissociation time of dCas9 was estimated to be ~200 min^48^, groups like R4, C2, and I2 exhibited apparently shorter interdivision cycles, suggesting that replication initiation did not halt when DnaA boxes were partially blocked. Second, the ordering of inhibitory effects was not entirely consistent with DnaA binding affinities of the target boxes, nor with results of a previous study^49^.

**Figure 2.**
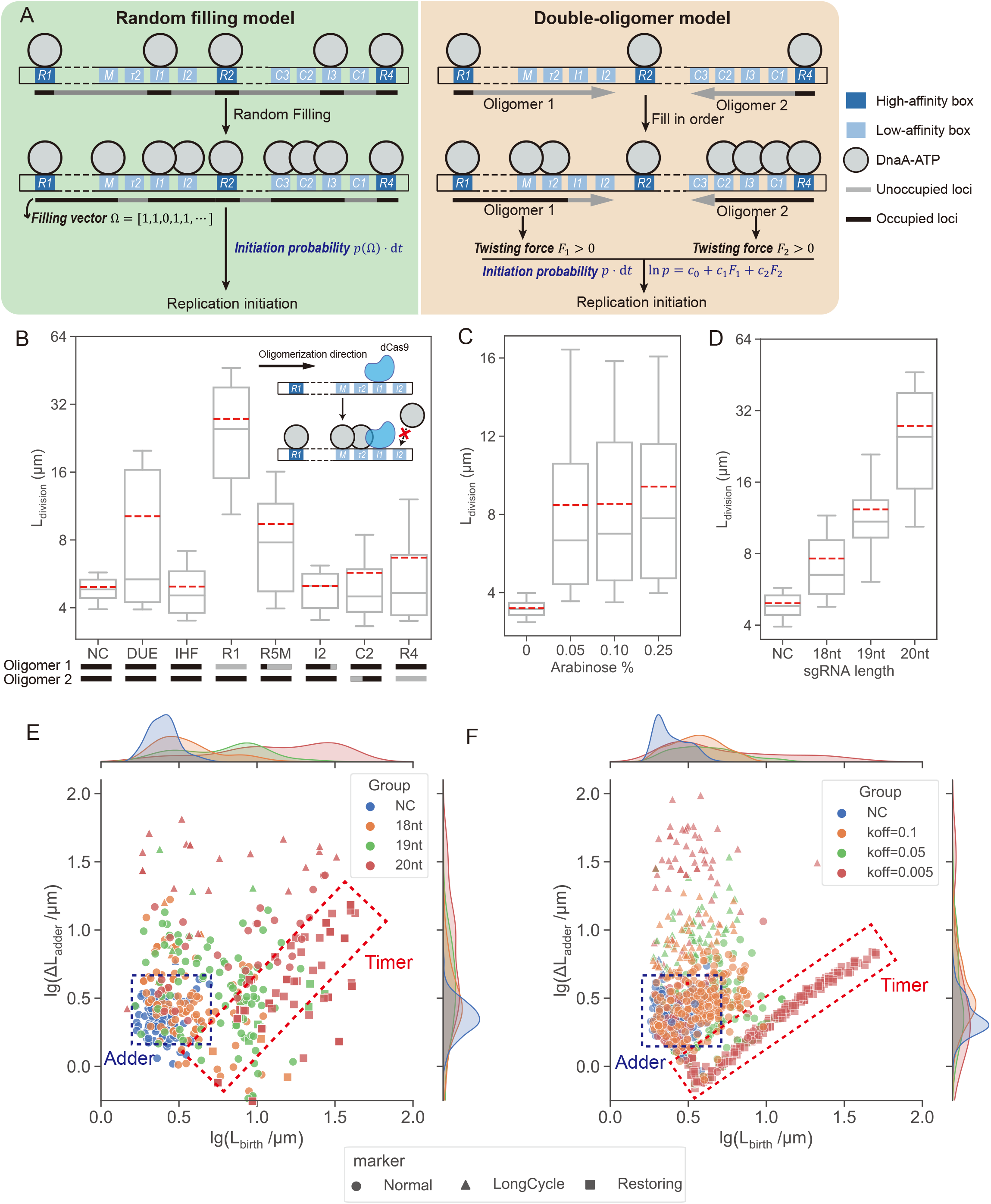
Tunable initiation perturbation by *CRISPRori* led to increased division lengths and the breakdown of the division adder. **A**. Two models for differential inhibition by *CRISPRori* targeting different DnaA boxes. Although the R1 box and the R5M box (M) are not adjacent on the linear DNA, IHF binds to the site in between and bends the DNA to bring both boxes together^16^. In the random filling model, the probability of replication initiation is a function of a filling vector in which 1 represents an occupied box and 0 an unoccupied box. In the double-oligomer model, the probability is assumed to depend on the twisting forces provided by two DnaA oligomers assembled on the left and right sides of the R2 box. **B-D**. Cell lengths before division for cells treated by *CRISPRori* targeting different sites in *oriC* (B), at different inducer concentrations (C), and with truncated sgRNAs (D). In B, the black bars below the *x*-axis indicate the maximal possible lengths of two oligomers with dCas9 occupying the box, and the grey bars show the full lengths. The inset shows a schematic of the occlusion effect of dCas9 to DnaA oligomer formation. In C and D, the targeted boxes were R5M and R1, respectively. In B and D, the inducer concentrations were all 0.25%. *N*_cell_ = 25~201. NC: negative control. For simulation results, see **Supplementary Fig. 4**. **E-F**. Experimental (E) and simulation (F) results showing the breakdown of the division adder under replication initiation perturbations. The increased cell length before division became widely distributed as perturbation strength increased. Filled triangles represent significantly elongated cells due to dCas9 inhibition, whereas filled squares were cell lineages rapidly dividing to restore to normal growth upon dCas9 release (see also **Fig. 3E**). In this phase, cells obeyed the timer principle, by which division happened as soon as a new replication cycle was completed (~15 min).

Combining with a DNA-centric cellular growth model that linked replication initiation to growth phenotypes (see next section), we tested two alternative models for replication initiation. The first “random filling” mechanism, where the probability of initiation depended on the number of boxes bound by DnaA in the *oriC* region **(Fig. 2A)** was rejected outright, since it predicted that cellular elongation scaled with the affinity of the targeted box to DnaA. Previous work showed that DnaA boxes are arranged on both sides of the R2 box in a head-to-head orientation. DnaA monomers bound to the strong binding sites R1 or R4 serve as points of nucleation to independently recruit other DnaA monomers onto low-affinity boxes to extend the left and right oligomers, both of which contribute to DNA unwinding^50^. Based on this, we subsequently hypothesized a two-site DnaA nucleation and oligomerization model **(Fig. 2A)**, by which the probability of initiation depended on the torsion provided by completely or partially assembled DnaA oligomers^41^. The location of the targeted box entered into the model by affecting the size of DnaA oligomers that can form. For example, if dCas9 blocked the I2 box, the R1 oligomer should still extend to the I1 box, leading to a reduced but non-zero probability of replication initiation. We found the observed inhibitory strengths well recapitulated by this model (see details in **Supplementary Note 3**).

Next, targeting the low-affinity R5M site, we varied the inducer concentration up to 0.25% arabinose and observed monotonically increasing division lengths with increasing dCas9 expression levels **(Fig. 2C, Supplementary Fig. 4C)**. Using a four-state competitive binding model between dCas9 and DnaA to the *oriC* sites, we estimated the time delay of replication initiation under various *CRISPRori* strengths **(Supplementary Note 4)**. The model predicted a transition of population behavior from a diffusion-limited bimodal phase to a dissociation-limited homogenous phase in single-cell division times and elongation morphologies, which were recapitulated in experiments (**Supplementary Fig. 5**). We also modulated the inhibition strength by changing the complementary length between the guide RNA and its target site, which was reported to introduce less noise than dCas9 expression modulation did^51^. For the R1 DnaA box, progressive truncations of sgRNAs by 1-nt resulted in significant decreases in division length **(Fig. 2D, Supplementary Fig. 4B)**.

How cells maintain their sizes in a narrow range has been investigated extensively in previous research^24,28,31–33,52–54^. Some bacteria, such as *E. coli* and *B. subtilis*, were hypothesized to control their sizes following the “adder” principle by adding a constant volume between birth and division. The adder was artificially broken by the synthetic expression of DnaA and FtsZ proteins in a previous study^24^. Noticing that an ongoing replication cycle could prevent cell division through nucleoid occlusion, we evaluated how the division adder responded to initiation perturbation. As expected, our data indicated that the division adder showed large variation until it broke down in the face of excessive replication postponement at strong inhibition levels **(Fig. 2E,F, Supplementary Fig. 6)**. Additionally, a previous study indicated the adder principle held true for multi-nucleoid filamentous *E. coli* cells that naturally occurred under stress conditions^29^. In our study of single-nucleoid filamentous cells with very low DNA-cytoplasm ratios, however, divisions transiently followed the “timer” principle before cell states were restored **(Fig. 2F, Fig. 3E)**.

**Figure 3.**
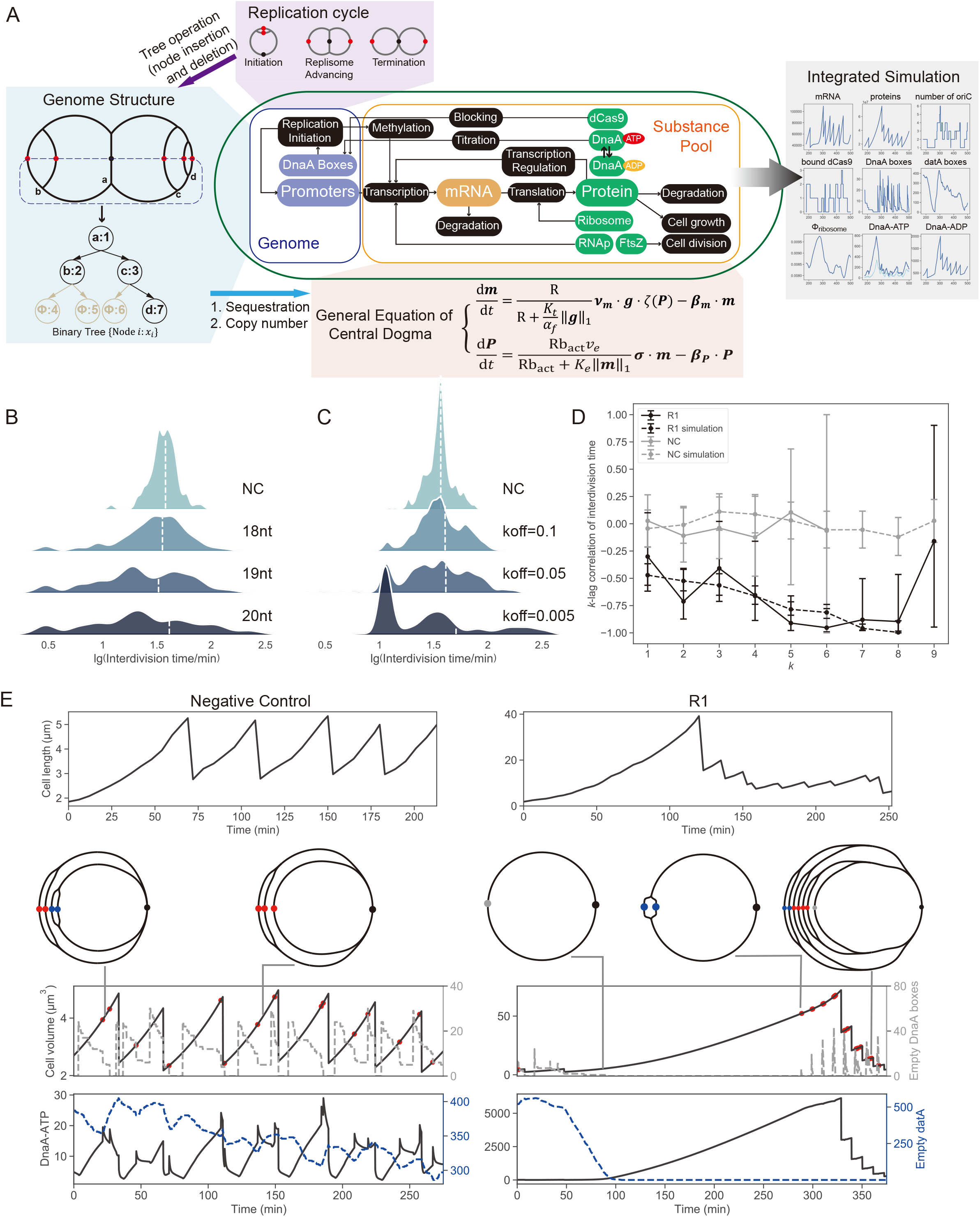
Cell division cycle shows homeostasis under initiation perturbations. **A**. Schematics of general bacteria growth model coupling protein expression, cell growth and cell cycles. **B-C**. Experimental (B) and simulated (C) distribution of the interdivision time under different replication inhibitory strengths, as tuned by the sgRNA length or the dCas9-sgRNA-DNA dissociation rate. The dashed lines marked the mean interdivision times. **D**. Correlation in interdivision times along cell lineages. The *k*-lag correlation is defined as the correlation between the current interdivision time (generation 0) and the mean interdivision time for generations 1 through *k*. In all lineages, only long interdivision cycles (*t*_d_>*t*_threshold_=60 min) were taken as generation 0. Error bar shows the 95% C.I. of the correlation coefficients. **E**. Experimental and simulated lineages for a normally replicating cell line (left) versus a *CRISPRori* inhibited cell line (right). The second row of cartoons shows the replication status of bacterial genomes at various time points, where black points show the *ter* site, blue points show SeqA sequestered *oriCs* that were accessible to neither dCas9 nor DnaA, gray points show dCas9-bound *oriC*s, and red points show the *oriC*s completely accessible to DnaA. The red dots in the lineage plots indicate replication initiation events. For more details of the simulated lineages, see **Supplementary Fig. 9** and **Supplementary Video 1&2**.

Taken together, these results showed that the *CRISPRori* synthetic system was highly tunable and quantitatively predictable in perturbing genomic replication initiation.

### A DNA-centric growth model for *Escherichia coli*

Over the past decade, bacterial growth has been modeled extensively as a consequence of the nutrition flux and dynamic cellular resource allocations^11,12,34–37,55^. Because the DNA content in bacterial cells has not been experimentally perturbed to large extents, these models largely neglect the contribution of genome dynamics in cell cycle progression.

Gene dosage effect has long been observed in prokaryotes, and recent studies have begun to fathom its underlying molecular mechanisms in relation to the partition of RNA polymerases (RNAP)^22,23,56^. In order to answer whether the cellular DNA content has an impact on general physiology, we proposed a mathematical model of cellular growth, taking DNA content as a central variable governing cell cycle progressing and general gene expression (**Fig. 3A**). The model was built on three major hypotheses:

1. RNAP and ribosomes were evenly distributed to all promoters and the ribosome binding sites on messenger RNAs, respectively.
2. For unregulated, constitutively expressed genes, firing frequencies of transcription and translation were dependent on the local concentration of RNAP and ribosomes, respectively, in a Hill-type manner.
3. DnaA governed the timing of DNA replication initiation, and cell division was contingent on FtsZ concentration and the presence of at least two nucleoids.

A detailed description of the model and the computation methods can be found in **Supplementary Notes 1 and 2.** Other notable processes previously reported to have affected DnaA-dependent replication initiation, including *oriC* sequestration, DnaA titration, auto-repression of DnaA transcription, the equilibrium between ATP-bound and ADP-bound form of DnaA, regulatory inactivation of DnaA (RIDA), etc.^16,25^, were all taken into consideration in the model. The criterion for DNA replication initiation as well as the model for dCas9-DnaA competitive binding were the same as described in previous sections. To avoid artifacts, most of the parameters were directly taken or estimated from previous measurements, with only the exponential growth rate and the free energy DnaA oligomers provides to unwind DNA fitted to the experimental results of the current study (**Supplementary Note 6**). Due to the stiffness of the system, a mixed method of Langevin equations and Monte Carlo sampling was used for numerical simulations, which enabled us to obtain high-resolution dynamic pictures of intracellular processes, where states of every replication origin (replicated and released, dCas9-bound or sequestered) were independently tracked (**Supplementary Note 5**).

As a control, we verified that at normal DNA-cytoplasm ratios, the model well recapitulated the quantitative phenomenological relations discovered and validated in past decades, including the Schaechter-Maaløe-Kjeldgaard’s exponential growth law^10^, Donachie’s hypothesis of initiation mass^30^, the adder principle^31–33^, and the linear relation between growth rate and ribosomal content as well as the unrelated protein expression^11^ (**Supplementary Fig. 7**). Besides, we asked whether the mechanisms we involved in the model are necessary for the cell to maintain normal physiology. We computationally “knocked out” RIDA, datA or DnaA auto-repression, and found that cells without any of these mechanisms exhibits either abnormal DNA replication or violation of “adder” principle (**Supplementary Fig. 14**). The simulation results show that our model is non-redundant and able to describe *E. coli* cell behavior in normal conditions, implying its potential in being extended to other regimes. In the following sections, combining experiment results with modeling, we focus on the cell cycle and cell growth at a DNA-cytoplasm ratio far below the normal range.

### Inter-division cycles exhibited homeostasis upon initiation perturbation

The breakdown of the division adder indicated severe cell cycle delays. Combining experimental observations and theoretical modeling, we first investigated how the temporary pause on genome replication caused by *CRISPRori* affected the cell’s division time. We collected interdivision times, i.e., the time interval between two consecutive cell division events, along single lineages of *CRISPRori* perturbed cells. Counterintuitively, even though cells were more likely to grow to larger sizes, the average interdivision time along their lineages remained unchanged (**Fig. 3B,C, Supplementary Fig. 8**).

A close examination of interdivision times revealed that delayed division did occur for elongated cells. The fraction of long cell cycles increased with perturbation strength, with the longest reaching ~300 min. Commensurately, extremely short division cycles also occurred more frequently, with the shortest being less than 3 min. This was reflected by the divergence of the interdivision time to a wide distribution under strongly perturbed conditions **(Fig. 3B,C)**. Interestingly, long cell cycles were almost always followed by several unusually short cell cycles until division time finally relaxed to the lineage mean, as revealed by the negative correlations in the interdivision time series that extended over eight generations after a prolonged cell cycle **(Fig. 3D)**. The strong coupling between long and short cell cycles suggested a prompt compensation mechanism to maintain cell cycle homeostasis despite constant perturbations to a key process in cellular growth.

Numerical simulations well reproduced experimental observations of cell cycle homeostasis. Simulated dynamics of DNA content, free DnaA-ATP, and FtsZ along continuously dividing cell lineages were tracked **(Fig. 3E, Supplementary Fig. 9**, **Supplementary Video 1,2)**. During the stoppage of DNA replication initiation, the cell continued expressing proteins, leading to the excessive accumulation of the initiator protein DnaA and the division protein FtsZ. After dCas9 dissociated, the excess DnaA and FtsZ promoted rapid rounds of replication initiation and cell division, resulting in unusually short cell cycles. These simulation results suggested the vertical passage of important cell cycle control proteins served a non-genetic “memory” that restored the cells to their normal growth states. Analytical estimation suggested the interdivision time converged to the long-term mean on an exponential time scale. In this “relaxation” period, however, the rate of convergence was initially faster than that obtained from the adder principle for normally growing cells **(Supplementary Note 7, Supplementary Fig. 10)**. We concluded that scaling between the DNA content and key cell cycle proteins enabled cells to self-repair abnormal DNA-cytoplasm ratios. As will be discussed below, the DNA-cytoplasm ratio can promote pronounced growth rate changes.

In addition, we separately verified that division ring organization and division site selection were largely unaffected by *CRISPRori* perturbation. They obeyed the law described in a previous study on filamentous *E. coli* of normal DNA-cytoplasm ratio^44^, despite an increased noise that occasionally led to extremely small, nucleoid-free daughter cells (**Supplementary Fig. 11**).

### Cell growth degenerated to linearity at low DNA-cytoplasm ratios

A recently published theory and an experimental study in yeast revealed a linear mode of cell growth upon DNA limitation^6,40^. Since *CRISPRori* produced cells with extremely low DNA-cytoplasm ratios, we experimentally measured their growth rates when *CRISPRori* was induced at different strengths. We found significant perturbation-dependent growth retardation in *CRISPRori* treated cells. Both single-cell instantaneous growth rates (based on cell length, **Fig. 4A, Supplementary Fig. 12B-F**) and bulk growth trajectories (as measured by the fraction area covered by cells in a microfluidic chamber, **Supplementary Fig. 12A**) exhibited a gradual transition from exponential to linear growth following the induction of *CRISPRori*. However, filamentous cells with normal DNA-cytoplasm ratios maintained exponential growth for more than 2 hours and up to ~100 μm in length until cell death^57^.

**Figure 4.**
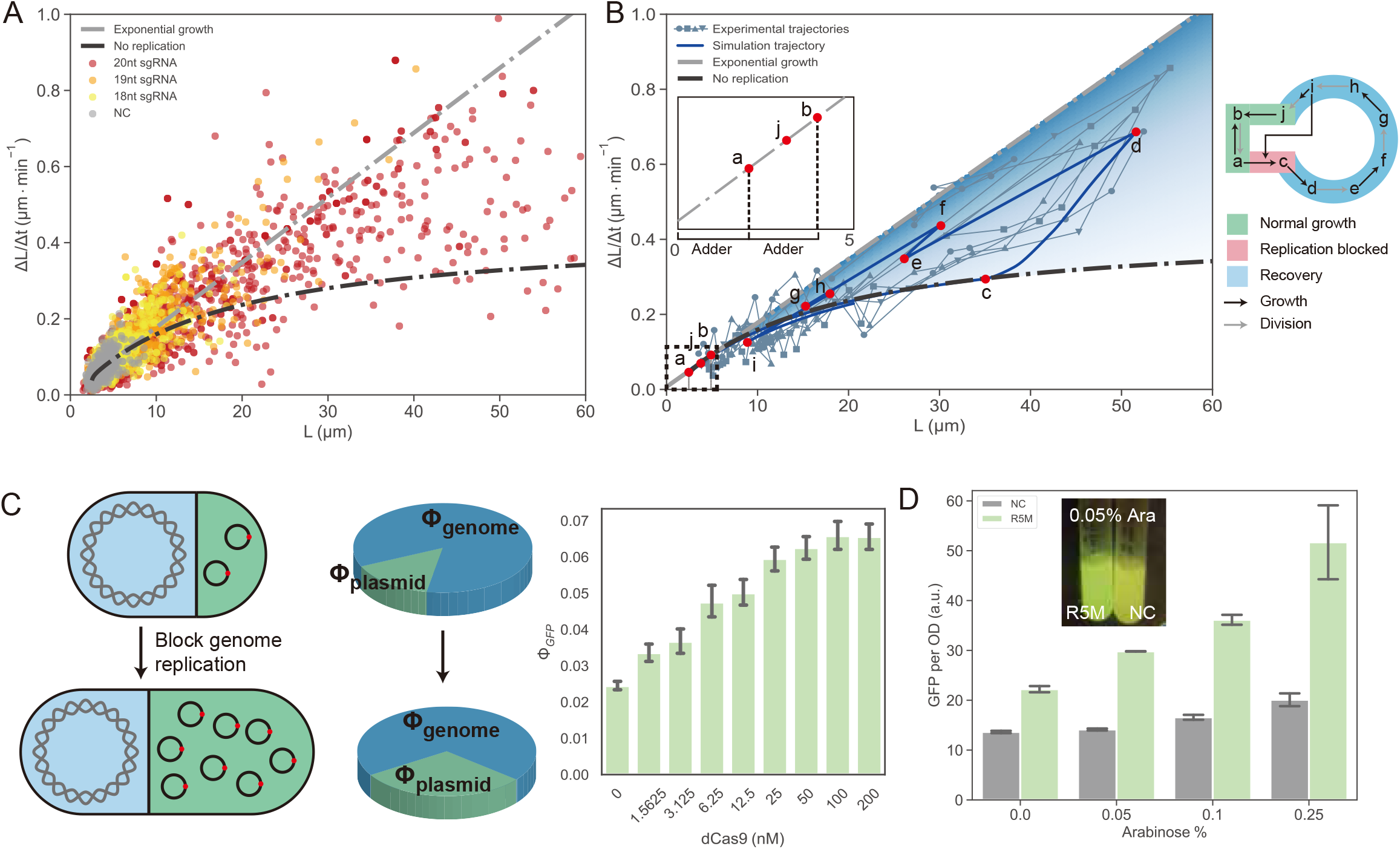
Transition from exponential to linear growth upon genome limitation enhanced expression of plasmid-encoded genes. **A.** Single-cell growth rate distributions. For exponential growth, the growth rate (ΔL/Δt) should be proportional to cell size L (gray dashed line). The slope of putative exponential growth decreased as perturbation strength increased. For severely perturbed cells (20bp sgRNA targeting R1), growth rate plateaued for elongated cells (black dashed line), indicating a linear growth pattern. **B.** Phase diagram for cellular growth states under a constant nutritional condition, as calculated by numerical simulations. The blue area between the exponential-growth line and the none-replication curve shows the permitted regime for balanced cell growth. Shading reflects the DNA:cytoplasmic ratio (darker shades for higher ratios). Trajectories of four experimentally tracked cell lineages (the R1 box blocked by dCas9, arabinose=0.25%), are plotted as gray curves with different markers. The thick blue curve shows a simulated trajectory, with letters indicating transitions between the normally replicating and the replication suppressed states as illustrated on the right. The inset is an enlarged view of the regime for non-perturbed cell states. **C.** Schematic (left) and simulated (right) intracellular resource allocation between the bacterial genome and the plasmids under normal and replication suppressed conditions. Φ denotes the proteomic fraction. Error bars show 95% C.I. of Φ_GFP_ for cells at 6 hours in simulation. **D.** Bulk measurements of the plasmid-borne GFP expression with genomic replication initiation inhibited by *CRISPRori*. Error bars show s.d. of three experimental replicates. For other measures of GFP expression, see **Supplementary Fig. 13**.

In the phase plane of cell size (*L*) and cell growth rate (d*L*/d*t*), the observed cell state at any instantaneous time fell within a “permitted growth” regime enclosed by a straight line for exponential growth and a curved line that leveled off at a constant linear growth rate **(Fig. 4B)**. The boundaries were fully mapped out by numerical simulations and were dependent on a constant nutrient supply. As indicated by the model, cells with high DNA-cytoplasm ratios (i.e., multi-nucleoid cells) always collapsed on the exponential boundary because the production of RNAP and ribosomal components were well balanced by the replicating genome. The permitted growth regime was expanded by decreasing nucleoid-cytoplasmic ratios, with which the accumulating RNAP and ribosomes gradually saturated the available gene copies and mRNAs, respectively. On the linear boundary, cellular production was strongly suppressed by a dearth of gene templates on the single genome copy. In agreement with theory, we found lineages unaffected by *CRISPRori* fluctuated around the exponential boundary in a narrow size range (normal size plus the adder), whereas *CRISPRori* treated lineages deviated toward the linear growth boundary, made large excursions into the permitted growth regime, and retreated to the exponential growth boundary in a few rounds of cell divisions **(Fig. 4B)**. These results underscored the essentiality of the balance between genome number and the cytoplasmic content in maintaining exponential growth, and the cells made their way back to normal through short division cycles.

### Limiting genomic DNA enhanced the expression of plasmid-encoded genes

The previous analysis suggested that gene copy numbers played a key role in intracellular resource allocation. In the presence of *CRISPRori*, as genes are mostly saturated by RNAP and mRNAs saturated by ribosomes, an intracellular resource surplus could be generated. We speculated that these spared resources might be redirected to plasmid-encoded genes for a high yield of recombinant proteins **(Fig. 4C)**.

We tested the expression of a plasmid-borne *gfp* gene in *CRISPRori* inhibited cells. The gross production, unit production (per OD600 unit), and single-cell fluorescence were measured under a series of arabinose concentrations. We found that replication perturbation indeed increased the unit GFP production with inducer concentration up to 0.25% **(Fig. 4D)**, while the gross production peaked early at mild perturbation strengths of 0.05-0.1% arabinose (measured by plate reader, **Supplementary Fig. 13A**). The non-monotonicity between gross production and perturbation strength was attributed to the competition between the increased GFP production rate and the dampened cellular growth rate. The single-cell fluorescence as measured by flow cytometry increased monotonously with arabinose concentration (**Supplementary Fig. 13B**).

Assuming plasmid replication depended on cytoplasmic DnaA concentration but was not inhibited by *CRISPRori*, model-simulated proteomic fraction of GFP behaved qualitatively the same with experimentally measured fluorescence **(Fig. 4C, Supplementary Fig. 13C)**. The agreement suggested that bio-production can be optimized through the redistribution of intracellular resources on DNA.

### *CRISPRori* enabled bidirectional tuning of the plasmid copy number

In theory, *CRISPRori* can be applied to any genetic system with a sequence-specific replication initiation mechanism. In particular, plasmids use diverse mechanisms to precisely regulate their copy number, which also mainly occur at the origin of replication^58^. Unlike bacterial chromosomes, many plasmids typically use antisense RNAs (or both RNAs and proteins) as a regulatory strategy^59^. For example, in the p15A origin, two promoters with opposite orientations direct the synthesis of a long RNA (RNA II) as the replication primer and of a short RNA (RNA I) that forms a duplex with the primer to block replication initiation^60^.

We tested the ability of *CRISPRori* to modulate plasmid replication both ways. The *CRISPRori* system was placed on a low copy number plasmid of the pSC101 origin. A plasmid containing a p15A origin and a constitutively expressed *mrfp* gene was used as the target. Two sgRNAs were designed to target RNA I and the initiation site, respectively **(Fig. 5A)**. The RFP fluorescence was measured to quantify the plasmid copy number^61^. In a concentration-dependent manner, fluorescence decreased when the initiation site was targeted but increased when the replication inhibitor RNA I was targeted. Collectively, bidirectional replication modulation by *CRISPRori* led to from 1/10 to 3-fold plasmid copy number variation at 0.25% arabinose concentration **(Fig. 5B)**. Guided by theory, flexible control of chromosome and plasmid DNA contents by *CRISPRori* might provide a novel way of adjusting intracellular resource allocation for the optimization of bio-production.

**Figure 5.**
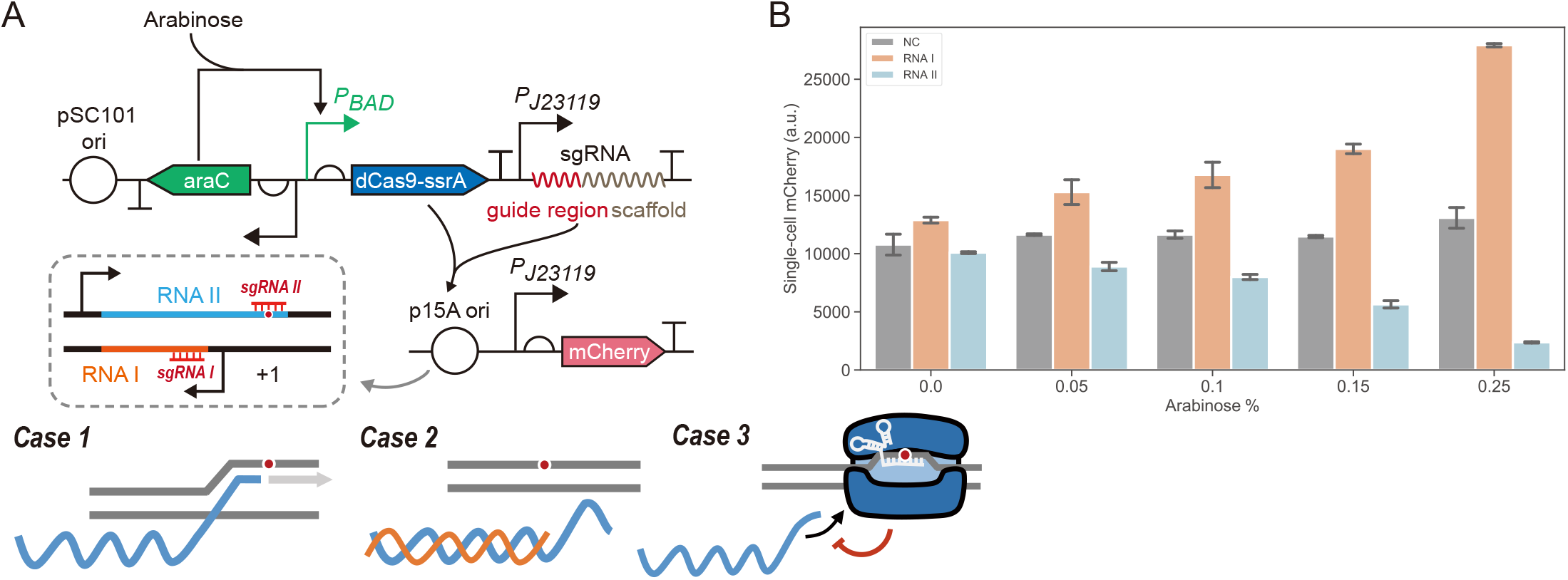
Bidirectional control of p15A plasmid copy number using *CRISPRori*. **A.** Copy number control of p15A plasmid by antisense RNAs and *CRISPRori*. The *CRISPRori* system was placed on a plasmid of pSC101 origin. The p15A plasmid harbored a constitutively expressed *mrfp* to indicate its copy number. At the p15A origin, RNA II (processed by RNaseH, not shown) acts as a primer for DNA polymerase to initiate DNA replication at the +1 site (Case 1). RNA I can hybridize with RNA II, preventing DNA-RNA II hybrid formation and replication initiation (Case 2). Two sgRNAs were designed to down-regulate RNA I and block the initiation site (Case 3), respectively. **B.** Copy number changes under the influence of *CRISPRori*. When targeting RNA I (sgRNA I) or the initiation site (sgRNA II), plasmid copy number was up- or down-regulated, respectively. Error bars show s.d. of three experimental replicates.

## Discussion

In this study, we applied dCas9-based DNA replication control to investigate how DNA replication initiation is coupled to cell division and cell growth. Other ways of halting replication include initiation sequence deletions and knockdown of components of the replication machinery. However, they suffered from lethality, pleiotropic effects, or functional redundancy^62–64^. A few previous studies used similar dCas9 systems to regulate replication initiation both *in vivo* and *in vitro*^49,65,66^. Our design of *CRISPRori* was more compact and optimized. And we showed *CRISPRori* provides tunable and reversible ways to regulate replication initiation.

The system allowed modeling that quantitatively elucidated the molecular processes underlying DnaA-mediated replication initiation and the competitive binding between dCas9 and DnaA. For example, the diverse effects when targeting different *oriC* boxes were explained by a double oligomer model, which showed consistency with a previous observation that nearly the entire right half of *oriC* was dispensable for proper functioning^50^. With almost all parameter values reported in literature, these mechanistic models coupling to a transcription-translation-based growth model faithfully recapitulated phenotypes induced by *CRISPRori*, including those related to cell cycle homeostasis and growth transitions. They together function as a multiscale model with sufficient molecular details that could serve as a foundation for other cellular physiological models. The model was solved through Langevin dynamics, which left out sources of molecular noise in, e.g., transcription and translation processes. How these noise properties finetune the population growth behavior remains a meaningful question to pursue.

By producing extreme DNA-cytoplasm ratios in live cells, *CRISPRori* enabled deep exploration into bacterial physiological regulation. One important discovery is the homeostasis of the division cycle under replication initiation perturbation. A previous study found that the division adder was independent of initiation control^24^. However, perturbation applied in the study was through oscillating DnaA, which was mild enough to not have changed the division size as reported previously^63^. In comparison, the *CRISPRori* experiments significantly affected division size and broke the division adder, fundamentally violating the *τ_cyc_* = *C* + *D, D* > 0 condition^24^. We showed that even in such conditions, the homeostasis of division cycles was maintained along single lineages. Further, we found the timer mechanism applied during the relaxation from a replication-inhibited state to the steady growth states, driven by the completion of new rounds of DNA replications. This represented a far-from-steady-state response. The exponential convergence rate for the timer mechanism was higher than that of the adder mechanism, which usually applies to noise-driven cell cycle heterogeneities near the steady state.

Another important discovery is the DNA-centric growth law, i.e., scaling between DNA and cytoplasmic proteins tuned cellular growth between two growth modalities. Previously, growth rate was recognized as a nutrition-dependent parameter and found to be internally constrained by the allocation of intracellular resources to ribosome synthesis^11^. Here, we demonstrated that, as DNA concentration decreased, widely accepted exponential growth degenerated into linear growth. Instead of ribosomes, DNA became the limiting factor governing cell growth^38,39^. This has been predicted by a theory before, but it is the first time that it was observed in live bacteria^40^.

Taken together, these observations suggested higher plasticity in DNA-cytoplasm ratios as well as growth modes for bacterial than for eukaryotic cells. One reason might be the relative independence between DNA replication and cell division, with which bacterial cells can rescue themselves by triggering consecutive divisions even at the cost of generating non-proliferable daughter cells. Unfortunately, eukaryotic cell cycle progression is controlled so tightly that division cannot occur without chromosome replication.

These fundamental findings inspired a potential application of the DNA-centric growth law and the *CRISPRori* system in bacterial physiological engineering. We showed *CRISPRori* regulation enabled production enhancement of recombinant proteins by directing spared RNAP and ribosomes to plasmid-encoded genes. It is also conceivable that energetic expenditure on DNA replication and cell division can be saved and used on product synthesis during replication initiation suppression. Further, we demonstrated bidirectional regulation of plasmid copy number by *CRISPRori*, implying its versatility for different biological systems, as we have accumulated massive sequencing data and annotations about the replication origin^67^ and proved Cas9’s compatibility with diverse host organisms^68^. The system may be used for balanced resource partitioning between host *vs*. synthetic genes, which provides a potentially better solution than unilaterally tuning up the plasmid copy number and letting synthetic functions exploit cellular resources. Lastly, the system’s reversibility can be leveraged to ensure the long-term robustness of engineered microbial cells.

## Material and Methods

### Strains and culture

*E. coli DH5α* strain was used for molecular cloning and measurements throughout this study. Incubations were carried out using a Digital Thermostatic Shaker (AOSHENG) at 37°C and 1,000rpm., using flat-bottom 96-well plates sealed with sealing film (Corning, BF-400-S). LB medium was used for molecular cloning and overnight incubation after picking single colonies from the plate. Before all measurements (DNA-cytoplasm ratio, growth monitoring in microfluidic chip, fluorescence), cells were cultured in M9 medium, according to a set of previously reported experimental procedures^69^ to ensure that gene expression was at the steady state. Antibiotic concentration used in this study was: Ampicillin (50 mg/mL), Chloramphenicol (17 mg/mL).

### Plasmid construction

The information on genetic parts used in this study can be found online (https://2019.igem.org/Team:Peking/Parts). The target sequence was inserted into sgRNA expression cassette using Golden Gate assembly (for more details, please refer to http://parts.igem.org/Part:BBa_K3081058). *CRISPRori* plasmids were constructed using pRG vector^70^, except in the plasmid copy number control experiment, where pSB4A5 vector was used. GFP-producing plasmid in expression enhancement experiment was constructed using pSB1C3 vector. The RFP-producing plasmid that indicated plasmid copy number was constructed using pSB3C5 plasmid.

### Monitoring cell growth in the microfluidic device

For detailed information about microfluidic chip design, fabrication, and operation, please refer to the original literature^46^. Before transferred into microfluidic chips, cells were inoculated from LB plate and cultured in LB medium overnight, then diluted into M9 medium and cultured for 3 hours. In the chip, cell growth was continuously monitored for 10 hours by taking pictures with a time interval of 3 min. For each group or treatment, 8 microchambers were selected as parallels.

### Nucleoid staining and observation

DAPI was used to stain the nucleoids in *E. coli* with a working concentration of 10 μg/mL. About 15 minutes after mixing bacteria with DAPI, the medium was replaced by PBS through precipitation-resuspension. After washing three times, the bacteria were available for microscopic imaging. Nucleoids were observed under laser scanning confocal microscope (UltraVIEW VoX, PerkinElmer, Inc.). Z-axis scanning for 2 μm with 0.2 μm each step overcame the imaging difficulty caused by rising and fall along the long cell body.

### Cell cytometry and plate reader

Single-cell fluorescence was measured using the flow cytometer (BD LSRFortessa) with appropriate voltage settings. For each assay, at least 20,000 events were collected. Flow cytometry data were analyzed using FlowJo software to obtain the arithmetic average of fluorescence intensity. For batch culture, the microplate reader (Thermo Scientific Varioskan Flash) was used to measure the fluorescence intensity and OD value. For both approaches, each group had at least 3 parallels.

### Image analysis

Volocity Demo 6.1.1. was used for confocal microscopy image analysis. To identify all nucleoids in one cell, XY planes were merged first; then contrast enhancement was performed. ImageJ software was used for cell dimension analysis (length and width), which was acquired by manually tracing cell lineages and doing measurements (using “segmented line” or “straight line”) at each time point. Time-lapsed cell length data can be further analyzed to calculate interdivision time and instantaneous growth rate. The coverage area was automatically calculated by creating an ImageJ Macro script.

### Data analysis and numerical simulations

All data analysis pipelines and numerical simulations are performed with MATLAB R2020a and Jupyter lab (Python 3.8). For modeling and simulation details, see

## Supplementary Notes 1-6

## Acknowledgments

We sincerely thank members of 2019 Peking University iGEM (International Genetically Engineered Machine Competition) team for doing preliminary works. We thank the Computing Platform of the Center for Life Science. We thank Fengyu Zhang and Prof. Chunxiong Luo for providing us with microfluidic chips and technical supports; Zhiwen Zhang for providing us with technical supports on microscopy; Prof. Qingsong Wang, Prof. Xinqiang He for lab equipment supports; Prof. Yihan Lin, Yihao Zhang and other colleagues for invaluable discussions. This work was supported by the National Key R&D Program of China (No.2020YFA0906900), the Natural Science Foundation of China (No. 31901063, No. 12090054) and the Top Notch Project 2.0 of PKU School of Life Sciences.

## Author contributions

S.Z., B.L., L.Q., P.W., Q.O., C.L. conceived the project. S.Z., B.L., L.Z., X.Q., and Y.S. designed and performed experiments. B.L., S.Z., L.Z. analyzed experimental data. B.L. and L.Q. constructed mathematical models and did numerical simulations. B.L., S.Z., and L.Q. created figures and wrote the manuscript.

