## Supplementary Notes and Figures for "Scaling between DNA and cell size governs bacterial growth homeostasis and resource allocation"

### A general theoretical framework coupling DNA replication, cell division and cell growth

#### Supplementary Note 1. Mathematical and computational descriptions of gene copy numbers on the dynamic genome

i. Gene copy numbers over a cell cycle

In this section, we show that DNA content can influence multiple cellular events, including global protein expression and cell cycle events. Following the idiomatic terminology, these influences are known as the gene-dosage effect.

Before giving a detailed description of how these effects actually are implemented, we would like to demonstrate how the gene copy number changes over the cell cycle and how these changes result in variant average gene copy numbers. We represent the location of a certain gene $i$ as $L_i \in [0,1]$, where the replication origin is at 0 and the terminus is at 1. Two replisomes form after a burst of DNA replication, and both of them are assumed to be symmetric with respect to the replication origin. If there are currently $n$ replication cycles on the genome with time-dependent replisome locations $\boldsymbol{x}(t) = [x_1, x_2, x_3, \cdots, x_n]$ $(x_m > x_{m+1}, m = 1,2,\cdots,n-1)$, the genomic fraction of gene $i$ located at $x < L_i \leq x_k$ is

$$\psi_{i,x(t)} = \frac{g_{i,x(t)}}{\sum_j g_{j,x(t)}} = \frac{2^k}{\sum_m 2^m G_{(x_{m+1},x_m]}} \qquad \textit{Eq. 1.1}$$

where $g_i$ is gene $i$'s copy number and $G_{(a,b]}$ denotes the number of genes located within $(a,b]$. We let $x_0 = 1$ and $x_{n+1} = 0$. For simplicity, we assume that the inter-replication time is a constant $\Delta t_i$ and the time to complete the replication is a constant $C$. Therefore, there are at most $n_{max} = \left\lceil \frac{C}{\Delta t_i} \right\rceil$ replication cycles on the genome. For a gene whose $L_i \in \left(\frac{k\Delta t_i}{C}, 1 - \frac{(n_{max}-1)\Delta t_i}{C}\right]$ $(k = 0,1,\cdots,n_{max}-1)$, the average gene dosage follows

$$\overline{g}_i = \frac{2^{n_{max}-k-1}}{L}\left[L + \frac{(n_{max}-1)\Delta t_i}{C} - L_i\right] \qquad \textit{Eq. 1.2}$$

As for a gene whose $L_i \in \left(1-\frac{(n_{max}-1)\Delta t_i}{C}, \frac{(k+1)\Delta t_i}{C}\right], k=0,1,\cdots,n_{max}-1$, the average gene dosage follows

$$\overline{g}_i = 2^{n_{max}-k-1} \qquad \textit{Eq. 1.3}$$

For example, if there is a bacterial cell with a 40-min *C*-period and 25-min inter-replication period, the average dosage of the gene located at $L$=1/4 is 2.4, a gene located at $L$=1/2 is 2, and gene located at $L$=3/4 is merely 6/5.

It is more complicated to calculate the average genomic fraction, as the total DNA content is continuously changing over the DNA replication cycle, and it requires knowledge about the genomic distribution of all genes. Thereby, we first derive the time-dependent total gene copy number. For convenience, we treat all genes as points with zero-length and evenly distributed along the genome. We set the time start point at a replication initiation event. For $t \in [0, C-(n_{max}-1)\Delta t_i]$, the total gene copy number should be

$$g(t) = \sum_j g_j = G\left[1 + \frac{(2^{n_{max}}-1)t + (2^{n_{max}} - n_{max} - 1)\Delta t_i}{C}\right] \qquad \textit{Eq. 1.4}$$

where $G \equiv G_{[0,1]}$ and $G \approx 4000$ in *E. coli*. Similarly, for $t \in (C-(n_{max}-1)\Delta t_i, \Delta t_i]$:

$$g(t) = G\left[1 + \frac{(2^{n_{max}-1}-1)t + (2^{n_{max}-1} - n_{max})\Delta t_i}{C}\right] \qquad \textit{Eq. 1.5}$$

With the total gene copy number obtained, we can write the expression of the average genomic fraction as

$$\overline{\psi}_i = \frac{1}{\Delta t_i}\int_0^{\Delta t_i} \frac{g_i(t)}{g(t)} \mathrm{d}t \qquad \textit{Eq. 1.6}$$

From *Eq. 1.6*, we can get the analytical expression of $\overline{\psi}_i$. We do not give the complicated expression here, but raise an example as a case study for clarity. Again, if there is a bacterial cell with 40-min C-period and 25-min inter-replication period, the total gene copy number can be written as

$$g(t) = \begin{cases} \left(\frac{3}{40}t + \frac{13}{8}\right)G & 0 \le t < 15 \\ \left(\frac{1}{40}t + 1\right)G & 15 < t \le 25 \end{cases} \qquad \textit{Eq. 1.7}$$

Then the average genomic fractions of genes located at 1/4, 1/2, and 3/4 of the genome are $1.252/G$, $1.096/G,$ and $0.718/G$, respectively. We will later show that this genomic fraction directly relates to their transcription levels through RNAP allocation.

##### ii. The data structure of a bacterial genome

The genome of the bacterial cell can be modeled as a typical binary tree. Because of the advancing replisomes, bursts of DNA replication, and replication termination, the structure of the genomic binary tree is dynamically changing all the time. Given that, we define a genomic binary tree in the following way:

> **Definition 1.1** (Genome binary tree). A genome binary tree is an ordered tree data structure with $\boldsymbol{n}(\boldsymbol{n} \ge \mathbf{0})$ nodes, in which each node represents a pair of replisomes generated by the same DNA replication initiation event and has at most two children. Every node in the binary tree $(\boldsymbol{n} > \mathbf{0})$ is defined by its node number $\boldsymbol{i} = \mathbf{1}, \mathbf{2}, \cdots, \boldsymbol{n}$ and genomic location $\boldsymbol{x_i} \in [\mathbf{0}, \mathbf{1}]$.

A genome binary tree is supposed to have all the properties of regular binary trees. The node number is determined in the order from root to branch and from left to right. Specifically, the $k^{th}$ node should be located at the $(\mathrm{k} - 2^{\lfloor \log_2 k \rfloor} + 1)^{\mathrm{th}}$ position in the $(\lfloor \log_2 k + 1 \rfloor)^{\mathrm{th}}$ layer. Based on the biological facts and Definition 1.1, it is easy to get following theorems on the genome binary tree:

> **Theorem 1.1** (Order of replication fork). Given a genome binary tree and two nodes in this tree with node number $\boldsymbol{i}$ and $\boldsymbol{j}$, if $\boldsymbol{j} > \mathbf{2}\boldsymbol{i}$, $\boldsymbol{x_i} < \boldsymbol{x_j}$
>
> **Theorem 1.2** (Replication origins). Given a genome binary tree with total node number $\boldsymbol{n}$, the number of replication origins $\boldsymbol{N}$ on the genome follows

$$N = \begin{cases} 2n - \sum_{i=1}^{n} d_i & n > 0 \\ 1 & n = 0 \end{cases} \qquad \textit{Eq. 1.8}$$

where $\boldsymbol{d_i}$ denotes the number of children of node $\boldsymbol{i}$.

Actually, Theorem 1.2 indicates that replication origins are located at every "empty" child node. Moreover, it was reported that the bacterial cell protects the genome from over-initiation of DNA replication by a mechanism named sequestration. During the sequestration period, the SeqA proteins bind to hemimethylated oriC and prevent re-initiation. This thereby leads to another noticeable property of the genome binary tree, which imposes the restriction on the minimal distance between a parent node and its children

**Theorem 1.3** (OriC sequestration). Given a bacterial genome with replication cycle *C* and sequestration period $\boldsymbol{\alpha C}$ $(\boldsymbol{\alpha} < \mathbf{1})$, for two nodes $\boldsymbol{i}$ and $\boldsymbol{j}$ on its binary tree, if $\mathbf{2}\boldsymbol{i} \le \boldsymbol{j} < \mathbf{2}(\boldsymbol{i} + \mathbf{1})$, $\boldsymbol{x_i} - \boldsymbol{x_j} \ge \boldsymbol{\alpha}$.

Following **Theorem 1.3**, we further have

**Theorem 1.4** Given a bacterial genome with replication cycle *C* and sequestration period $\boldsymbol{\alpha C}$ $(\boldsymbol{\alpha} < \mathbf{1})$, it has a maximum number of pairs of replication forks $\mathbf{2}^{\lceil \frac{1}{\alpha} \rceil} - \mathbf{1}$ and a maximum number of replication origins $\mathbf{2}^{\lceil \frac{1}{\alpha} \rceil}$.

α has been reported to be slightly less than 1/3 in *E. coli* in normal conditions, which means every genome in *E. coli* can have at most 15 replisome pairs and 16 oriCs.

The last section discussed the analytical expression of genomic fraction for a specific gene on the genome with a constant replication cycle and inter-initiation time. We will later show that inter-initiation time can be determined independently and changing over different cycles, so it is essential to connect the genomic fraction with the genome binary tree computationally.

**Algorithm 1** Calculate DNA content from the genome binary tree

**Input:** $\{i\}$: all node number; $\{x_i\}$: node locations

**Output:** $G$: DNA content

**if** $\{i\} = \emptyset$ **then**

No replication cycles

$G \leftarrow 1$

**else**

**Initialize** $G \leftarrow 0$

**for** $j \in \{i\}$ **do**

**if** $j = 1$ **then**

From replication terminus to the first replisome

$G \leftarrow G + 1 - x_j$

**else**

From parent node to child node

$G \leftarrow G + x_{\lfloor \frac{j}{2} \rfloor} - x_j$

**if** $2j \notin \{i\}$ **then**

After the node on the last layer

$G \leftarrow G + x_j$

**end if**

**if** $2j + 1 \notin \{i\}$ **then**

After the node on the last layer

$G \leftarrow G + x_j$

**end if**

**end if**

**end for**

**end if**

**Algorithm 2** Calculate DNA content from the genome binary tree

**Input:** $\{i\}$: all node number; $\{x_i\}$: node locations; $L$: gene location

**Output:** $G_{\mathrm{L}}$: copy number of a gene located at $L$

**if** $\{i\} = \emptyset$ **then**
  $G_L \leftarrow 1$
**else**
  **Initialize** $G_L \leftarrow 0$
  **for** $j \in \{i\}$ **do**
    **if** $x_j < L$ **then**
      **if** $\mathrm{j} = 1$ **then**
        $G_L \leftarrow G_L + 1$
      **else**
        **if** $L_{\left\lfloor \frac{j}{2} \right\rfloor} > L$ **then**
          $G_L \leftarrow G_L + 1$
        **end if**
      **end if**
    **end if**
    **if** $2j \notin \{i\}$ **and** $L_j > L$ **then**
      $G_L \leftarrow G_L + 1$
    **end if**
    **if** $2j + 1 \notin \{i\}$ **and** $L_j > L$ **then**
      $G_L \leftarrow G_L + 1$
    **end if**
  **end for**
**end if**

Algorithms 1 and 2 do not exclude the situation that genes located around the replication origin are not capable of getting transcribed during the sequestration period. We will later show that genes in this region do not bind with RNA polymerases and do not participate in resource allocation.

So far, we have analyzed the structure of bacterial genomes at the static state. However, as the replisome keeps advancing during the replication cycle, $L_i$ is time-dependent. The advancing rate of replisomes can be treated as a constant in the long run, in spite of instantaneous moves and stops (i.e., $\frac{\mathrm{d}x_i(t)}{\mathrm{d}t} = \frac{1}{C} = const.$). When $x_i$ reaches 1, replication termination occurs, where a single nucleoid divides into two.

> **Definition 1.2** (Replication termination). For a given non-empty genome binary tree, replication termination corresponds to the division of the tree into the left child tree and right child tree of the root node, with the replisome location unchanged.

Replication initiation is another obvious process that alters the genome binary tree structure by inserting a node on the last layer.

> **Definition 1.3** (Replication initiation). For an empty genome binary tree, if DNA replication initiation occurs, add a root node to the empty tree with a replisome location equal to zero. For a given non-empty genome binary tree, if DNA replication initiation occurs at the left (right) branch of node $\boldsymbol{i}$ which has less than two children, add a new node to the tree with node number $\boldsymbol{2i}$ ($\boldsymbol{2i+1}$ for the right branch) and replisome location zero.

**Supplementary Note 2. A global DNA-content-dependent cell growth model**

i. Gene dosage-dependent gene expression

Transcriptional activity of constitutive promoters has been proposed to depend on the local concentration of free RNA polymerase holoenzymes in a Michaelis-Menten manner[1]. We here use $v_{t,i}$ to represent the firing frequency of transcription of gene $i$, which is related to the number of free RNAP holoenzymes distributed to gene $i$, $R_{f,i}$.

$$v_{t,i} = \frac{v_{m,i} R_{f,i}}{R_{f,i} + K_t} \qquad \textit{Eq. 2.1}$$

where $v_{m,i}$ denotes maximal firing frequency when $R_{f,i} \to \infty$ and $K_t$ equals to $R_{f,i}$ at half maximal firing frequency. It can be treated as the intrinsic property of the gene promoter. Three major states of intracellular RNAPs were directly observed by super-resolution microscopy: DNA specifically bound, non-specifically bound, and free RNAP[2,3]. These states are in fast equilibrium, suggesting that free RNAPs comprise a fixed proportion in total RNAPs.

$$\sum_j R_{f,j} = \alpha_f R \qquad \textit{Eq. 2.2}$$

where $\sum_j$ is to sum over all genes (i.e., promoters), $R$ is the total intracellular amount of RNAP and $\alpha_f$ denotes the fixed fraction of free RNAP. Next, to obtain the expression of transcription rate, we use $v_t$ and $l_i$ to represent the elongation rate of RNAP and length of gene $i$. One gene can hold a number of RNAPs simultaneously, and thus the transcription rate of gene $i$ should be the product of the number of RNAPs held by gene $i$ and the relative elongation rate $v_t/l$.

$$r_{t,i} = \frac{v_{t,i} l}{v_t} \cdot \frac{v_t}{l} = v_{t,i} \qquad \textit{Eq. 2.3}$$

*Eq. 2.3* suggests that the transcriptional activity is elongation rate-independent. This RNAP-dependent manner coincides with that transcription in normal conditions is DNA-excessive but RNAP-limiting[4,5]. Previous work also demonstrated the reduction of free RNAP and transcription per gene with increasing intracellular promoters[6], suggesting that the RNAPs should be appropriately distributed to all promoters. In spite of the genome structure, here we assume even distribution of RNAP to genes:

$$R_{f,i} = \frac{g_i}{\sum_j g_j} \cdot \alpha_f R = \alpha_f \psi_i R \qquad \textit{Eq. 2.4}$$

in which we use $\psi_i$ to the genomic fraction of gene $i$. Therefore, the transcription equation can be written as

$$r_{t,i} = \frac{v_{m,i} g_i \cdot \left(\frac{R}{g}\right)}{\frac{R}{g} + \frac{K_t}{\alpha_f}} \cdot \zeta(b_i) \qquad \textit{Eq. 2.5}$$

where $\zeta(b_i)$ is the regulation function of exogenous regulatory factors of gene $i$, for instance, lactose analog IPTG for Lac operon and intracellular (p)ppGpp for rRNA and RNAP transcription. Notably, this transcription equation introduces genomic fraction (i.e., $\psi_i$), which reflects the gene-dosage effect on protein expression level as observed in *E. coli* and *B. subtilis*. These works found a several-fold higher expression level of genes close to the replication origin than genes close to the replication terminus[7-10].

The translation is the central step of protein synthesis, whose activity can be altered by ribosome content or nutritional conditions. Notably, ribosomes can be classified into two types: active ribosomes that are directly engaged in translation and inactive ribosomes that are not, and thus the translational activity is written:

$$r_{e,i} = \frac{m_i v_e Rb_{a,i}}{l_i} \qquad \textit{Eq. 2.6}$$

where the letter representations are as follows:

| **Variables** | **Physical meaning** |
| --- | --- |
| $r_{e,i}$ | Translational rate of mRNA of gene $i$ |
| $v_e$ | Peptide synthesis elongation rate |
| $Rb_{a,i}$ | Active ribosomes engaged in the translation of mRNA of gene $i$ |
| $l_i$ | Length of mRNA of gene $i$ |

Similar to transcription, we treat the number of active ribosomes on mRNA $i$ as a Michaelis-Menten function of local ribosome concentration:

$$Rb_{a,i} = \frac{Rb_{m,i} Rb_i}{Rb_i + K_e} \qquad \textit{Eq. 2.7}$$

$Rb_{m,i}$ is the ribosome number held by mRNA $i$ with the maximal density $\sigma_i$, which is an intrinsic property of the sequence from the Shine-Dalgarno sequence to the start codon:

$$Rb_{m,i} = l_i \sigma_i \qquad \textit{Eq. 2.8}$$

Joining *Eq. 2.7* and *Eq. 2.8*, we have:

$$r_{e,i} = \sigma_i v_e m_i \frac{Rb_i}{Rb_i + K_e} \qquad \textit{Eq. 2.9}$$

Moreover, the decline of protein synthesis implemented by transcription inhibition indicates that mRNA copy number also plays an essential role in ribosome allocation. Given the fact that not all ribosomes are actively translating, we can rewrite $Rb_i$ as

$$Rb_i = \frac{1}{\sum_j m_j}(Rb - Rb') = \frac{Rb - Rb'}{m} \qquad \textit{Eq. 2.10}$$

Therefore, the translation rate of mRNA of gene $i$ can be written as

$$r_{e,i} = \sigma_i v_e m_i \frac{\frac{Rb - Rb'}{m}}{\frac{Rb - Rb'}{m} + K_e} \qquad \textit{Eq. 2.11}$$

It has been reported that the elongation rate of the ribosome is growth rate-dependent[11,12], i.e. $v_e \equiv v_e(\lambda)$. As time for adding one amino acid to the peptide can be divided into the time for dehydration synthesis and for component diffusion, including enzyme EF-G, tRNA ternary complex (comprising aminoacyl-tRNA, elongation factor Tu (EF-Tu) and guanosine triphosphate (GTP)) and etc., its formulation can be specified as[11,12]

$$v_e = \frac{1}{\langle \tau_e \rangle} = \frac{1}{\frac{1}{k_{syn}} + \sum_c \frac{1}{ck_{on,c}}} \qquad \textit{Eq. 2.12}$$

ii. A general equation of protein synthesis

The translation equation indicates that the production rate of protein $i$ is

$$\frac{\mathrm{d}p_i}{\mathrm{d}t} = \sigma_i v_e m_i \frac{\frac{Rb - Rb'}{m}}{\frac{Rb - Rb'}{m} + K_e} - \frac{p_i}{\tau_{p,i}} \qquad \textit{Eq. 2.13}$$

where $\tau_{p,i}$ is the degradation time for protein $i$, and $m_i$ should be solved from

$$\frac{\mathrm{d}m_i}{\mathrm{d}t} = \frac{v_{m,i} g_i \frac{R}{g}}{\frac{R}{g} + \frac{K_t}{\alpha_f}} \cdot \zeta(b_i) - \frac{m_i}{\tau_{m,i}} \qquad \textit{Eq. 2.14}$$

For some genes whose transcription level is regulated by other proteins, such as transcription activators or repressors, the term $\zeta(b_i)$ may involve $p_j$ (maybe more than one). This ends up with a combination of $2G$ equations with $2G$ unknowns.

$$\frac{\mathrm{d}\boldsymbol{P}}{\mathrm{d}t}=\frac{(Rb-Rb')v_e}{Rb-Rb'+K_e\|\boldsymbol{M}\|_1}\boldsymbol{\Sigma}\cdot*\boldsymbol{M}-\boldsymbol{\beta_P}\cdot*\boldsymbol{P} \qquad \textit{Eq. 2.15}$$

$$\frac{\mathrm{d}\boldsymbol{M}}{\mathrm{d}t}=\frac{R}{R+\frac{K_t}{\alpha_f}\|\boldsymbol{G}\|_1}\boldsymbol{N_m}\cdot*\boldsymbol{G}\cdot*\zeta(\boldsymbol{P})-\boldsymbol{\beta_M}\cdot*\boldsymbol{M} \qquad \textit{Eq. 2.16}$$

A significant problem in solving *Eq. 2.15* and *Eq. 2.16* is to get transient values for ribosomes and RNAPs. Therefore, given that synthesis of r-proteins is the rate-limiting step of ribosome self-reproduction, it is proper to treat ribosomes as proteins simply. Then $Rb$ and $R$ can be solved by combining the following equations:

$$\begin{cases}\dfrac{\mathrm{d}Rb}{\mathrm{d}t}=\sigma_{Rb}v_e m_{Rb}\dfrac{Rb-Rb'}{Rb-Rb'+mK_e}\\ \dfrac{\mathrm{d}R}{\mathrm{d}t}=\sigma_R v_e m_R\dfrac{Rb-Rb'}{Rb-Rb'+mK_e}\\ \dfrac{\mathrm{d}m_{Rb}}{\mathrm{d}t}=\dfrac{v_{m,Rb}g_{Rb}\alpha_f R}{\alpha_f R+gK_t}\zeta(b_{Rb})-\dfrac{m_{Rb}}{\tau_{m,Rb}}\\ \dfrac{\mathrm{d}m_R}{\mathrm{d}t}=\dfrac{v_{m,R}g_R\alpha_f R}{\alpha_f R+gK_t}\zeta(b_R)-\dfrac{m_R}{\tau_{m,R}}\end{cases} \qquad \textit{Eq. 2.17}$$

iii. Balanced growth

Theoretically, the general protein synthesis equation solution can result in an accurate definition of cell growth. However, obtaining an analytical solution requires proper approximations.

As the transcription regulation function of very few genes can be specified, we coarsely classify all genes into two groups: genes whose proteomic fraction is a constant (i.e., $\phi_i=const.$) and genes with constitutive promoters (i.e., $\zeta(b_i)=1$). Here we address two limiting cases in which the equation can be linearized.

**Case 1:** $\alpha_f R\ll K_t g$ and $Rb\ll mK_t$.

This is the case when RNAPs are far from saturating the genome and ribosomes are far from saturating mRNAs, similar to standard growth conditions. *Eq. 2.14* can be written as

$$\frac{\mathrm{d}m_i(t)}{\mathrm{d}t}=\frac{v_{m,i}\psi_i(t)\alpha_f R(t)}{K_t}\cdot\zeta\big(b_i(t)\big)-\frac{m_i(t)}{\tau_{m,i}} \qquad \textit{Eq. 2.18}$$

We apply the approximation to *Eq. 2.18* that $\psi_i(t) \approx \overline{\psi}_i = \lim_{\tau\to\infty} \frac{1}{\tau}\int_0^\tau \psi_i(t)\mathrm{d}t$ and $\zeta(b_i(t)) \approx \zeta(\overline{b}_i)$. Therefore, solving *Eq. 2.18*, we have

$$m_i = C_i e^{-t/\tau_{m,i}} + \frac{\alpha_f v_{m,i}}{K_t}\overline{\psi}_i \zeta(b_i) e^{-t/\tau_{m,i}} \int R(t) e^{t/\tau_{m,i}} \mathrm{d}t \qquad \textit{Eq. 2.19}$$

Let $t \to \infty$, and we have

$$m_i^\infty = \frac{\alpha_f v_{m,i}}{K_t}\overline{\psi}_i \zeta(b_i) \lim_{t\to\infty} e^{-\frac{t}{\tau_{m,i}}} \int R(t) e^{\frac{t}{\tau_{m,i}}} \mathrm{d}t \qquad \textit{Eq. 2.20}$$

where $m_i^\infty$ is the steady-state value of $m_i$ if the integration term converges. For simplicity, we treat the degradation time for all mRNAs as equal, i.e., $\tau_{\mathrm{m,i}} \equiv \tau_m$. In other words, if time is long enough:

$$\mu_i^* \propto \frac{v_{m,i}\overline{\psi}_i \zeta(b_i)}{\sum_j v_{m,j}\overline{\psi}_j \zeta(b_j)} = const. \qquad \textit{Eq. 2.21}$$

Substituting into *Eq. 2.13* and applying the assumption that $Rb \ll mK_t$, we have

$$\frac{\mathrm{d}Rb}{\mathrm{d}t} = \frac{\sigma_{Rb} v_e \mu_{Rb}^*}{K_e}(Rb - Rb') \qquad \textit{Eq. 2.22}$$

Notably, the fixed portion of inactive ribosomes scales with total proteins, i.e., $\phi'_{Rb} = \mathrm{const.}$ Next, we turn to the total protein synthesis:

$$\frac{\mathrm{d}}{\mathrm{d}t}\sum_j p_j = \sum_j \frac{\sigma_j v_e \mu_j^*}{K_e}(Rb - Rb') = \frac{\langle\sigma\rangle v_e}{K_e}(\phi_{Rb} - \phi'_{Rb}) \qquad \textit{Eq. 2.23}$$

where $\langle\sigma\rangle = \frac{\sum_j \sigma_j \mu_j^*}{\sum_j \mu_j^*}$. Hence *Eq. 2.23* can be written as

$$\frac{\mathrm{d}}{\mathrm{d}t}\ln\sum_j p_j = \frac{\langle\sigma\rangle v_e}{K_e}(\phi_{Rb} - \phi'_{Rb}) \qquad \textit{Eq. 2.24}$$

Therefore, cell growth largely depends on the proteomic fraction of ribosomes:

$$\frac{\mathrm{d}}{\mathrm{d}t}\phi_{Rb} = \frac{\mathrm{d}}{\mathrm{d}t}\frac{Rb}{\sum_j p_j} = \frac{v_e}{K_e}(\phi_{Rb} - \phi'_{Rb})(\sigma_{Rb}\mu_{Rb}^* - \langle\sigma\rangle\phi_{Rb}) \qquad \textit{Eq. 2.25}$$

Let $\frac{\mathrm{d}\phi_{Rb}}{\mathrm{d}t} = 0$. We then have the steady-state solution:

$$\phi_{Rb}^* = \frac{\sigma_{Rb}\mu_{Rb}^*}{\langle\sigma\rangle} \qquad \textit{Eq. 2.26}$$

Substituting *Eq. 2.26* into *Eq. 2.24*, we then find that the steady-state changing rate of total proteins is a constant:

$$\frac{\mathrm{d}}{\mathrm{d}t}\ln\sum_j p_j = \frac{\sigma_{Rb}v_e\mu_{Rb}^*}{K_e} - \frac{\langle\sigma\rangle v_e}{K_e}\phi_{Rb}' = \text{const.} \qquad \textit{Eq. 2.27}$$

*Eq. 2.27* is the well-known exponential growth of bacteria. To avoid ambiguity, we define *Eq. 2.27* as the exponential growth rate, i.e., $\lambda_{exp}^* = \frac{\mathrm{d}}{\mathrm{d}t} ln \sum_j p_j$. We now return to the proteomic fraction of ribosomes by substituting *Eq. 2.27* into *Eq. 2.26*:

$$\phi_{Rb}^* = \phi_{Rb'} + \frac{K_e}{v_e\langle\sigma\rangle}\cdot\lambda_{exp}^* \qquad \textit{Eq. 2.28}$$

**Case 2:** $\alpha_f R \gg K_t g$ and $Rb \gg mK_t$.

This is when genome and mRNAs have been almost saturated by RNAPs and ribosomes, respectively, which describes a bacterial cell with a very low DNA-to-cytoplasm ratio. With the similar derivation process in **Case 1**, the mRNA equation can be written as:

$$\frac{\mathrm{d}m_i}{\mathrm{d}t} = v_{m,i}g_i\zeta(b_i) - \frac{m_i}{\tau_{m,i}} \qquad \textit{Eq. 2.29}$$

whose solution is

$$m_i = v_{m,i}g_i\zeta(b_i)\tau_{m,i} - C_i\tau_{m,i}e^{-\frac{t}{\tau_{m,i}}} \qquad \textit{Eq. 2.30}$$

Moreover, the translation equation is

$$\frac{\mathrm{d}p_i}{\mathrm{d}t} = \sigma_i v_e m_i = \sigma_i v_e\left(v_{m,i}g_i\zeta(b_i)\tau_{m,i} - C_i\tau_{m,i}e^{-\frac{t}{\tau_{m,i}}}\right) \qquad \textit{Eq. 2.31}$$

Solving *Eq. 2.31*, we have

$$p_i = \sigma_i v_e\left(v_{m,i}g_i\zeta(b_i)\tau_{m,i}t + C_i\tau_{m,i}^2 e^{-\frac{t}{\tau_{m,i}}}\right) + p_0 \qquad \textit{Eq. 2.32}$$

When the time is long enough, *Eq. 2.32* can be rewritten as

$$p_i = \sigma_i v_e v_{m,i}g_i\zeta(b_i)\tau_{m,i}t + p_0 \qquad \textit{Eq. 2.33}$$

which results in the linear growth of the cell. Here we define the linear growth rate as

$$\lambda_{lin}^* = \frac{\mathrm{d}V}{\mathrm{d}t} = \frac{v_e}{c_P^*}\sum_j g_j\sigma_j v_{m,j}\tau_{m,j}\tau_{m,j}\zeta(b_j) \qquad \textit{Eq. 2.34}$$

iv. A DNA-centric growth law

To find the relation between growth rate and DNA content, we define a central variable $X = \frac{\sum_j g_j}{\sum_j p_j}$ which is the ratio between the total gene copy number and the protein amount. Although *Eq. 2.17* is not directly mathematically tractable, we can apply several approximations to give a closed-form solution that provides us with physical intuitions. First, we assume a "reinforced balanced growth", where $\psi_{Rb}, \phi_{Rb}$ and $\phi_R$ are constants with a given $X$. We name it "reinforced" because this kind of balanced growth might not be dynamically reachable. Namely, we hypothesize that there is a bizarre cell whose gene copy number keeps scaling with its size, and its growth will eventually reach balance, but apparently, it is impossible in reality. Secondly, we assume the gene copy number $g(t) \approx \overline{g} = \lim_{\tau\to\infty} \frac{1}{\tau}\int_0^\tau g(t)dt = \text{const.}$

$$m_{Rb}^* = \frac{v_{m,Rb}\overline{g_{Rb}}\tau_{m,Rb}}{1 + \frac{K_t}{\alpha_f \phi_R} \cdot X} \qquad \textit{Eq. 2.35}$$

We define a general expression of balanced growth rate as $\lambda^* = \frac{1}{Rb^*}\frac{dRb^*}{dt}$. Substituting *Eq. 2.35* and *Eq. 2.17* into this expression, we eventually have

$$\lambda^* = \frac{v_e \sigma_{Rb} \psi_{Rb}^*}{\frac{\phi_{Rb}^*}{v_{m,Rb}\tau_{m,Rb}}\left(X + \frac{K_t}{\alpha_f \phi_R}\right) + K_e} \qquad \textit{Eq. 2.36}$$

#### Supplementary Note 3. Modeling replication initiation

i. Overview of DnaA-mediated DNA replication initiation

Accumulating evidence has been discovered for DnaA as the DNA replication initiator in *E. coli*[13-15]. As a highly conserved protein in bacterial species, DnaA can form a stable complex with both ATP and ADP, but only ATP-bound DnaA can efficiently initiate replication. Initiation of DNA replication has the most direct relation to intracellular DNA content while it is tightly regulated by DNA-dependent processes through multiple mechanisms. Modeling replication initiation can be divided into two parts, the regulation part, which discusses regulatory mechanisms on DnaA functioning through regions outside *oriC*, and the trigger part, which discusses the criterion for triggering initiation. In the latter part, we would elaborate on how DnaA boxes are distinct from each other.

There are several known DnaA-boxes in oriC to which DnaA can bind in a cooperative manner and take on a spiral shape[16,17]. DnaA shows variant affinities to different boxes, but notably, binding to low-affinity boxes, including R5M, τ2, I1, I2, C1, I3, C2, and C3, is crucial for complex activation and the unwinding of the AT-rich region[14]. In addition to DnaA-boxes in the oriC region, there are other DnaA-binding regions on the chromosome known as *datA* (DnaA titration) loci, known to be hundreds of consensus DnaA boxes on the *E. coli* chromosome based on biochemical analysis of purified DnaA proteins[18]. The existence of these boxes located outside oriC can have a remarkable influence on replication initiation by efficiently decreasing the free DnaA proteins in the cell.

Multiple other regulations on the DnaA-dependent DNA replication initiation exist in bacterial cells to ensure the precise timing of initiation and avoid re-initiation in the same replication cycle. As for the expression of DnaA itself, two fairly strong promoters govern the transcription of *dnaA* genes. The DnaA promoter region carries sequences homologous to DnaA-boxes, such that both promoters are negatively regulated by DnaA itself, but *in vivo* experiments show that this autoregulation effect is relatively weak[19-21]. *dnaA* transcription has been reported to be dependent on the growth rate[21-23].

Besides, the coding sequence of *dnaA* has a GUG start codon and a relatively poor ribosome binding site, and therefore translation efficiency of DnaA protein is merely about one-third of that for genes with an AUG start codon[24]. It is noteworthy that the DnaA protein itself is stable[25]. Early experiments have suggested a growth rate-dependent DnaA mRNA level but a growth rate-independent DnaA intracellular concentration[21,22,26], though the latter conclusion conflicts with a recent work in which DnaA concentration shows a non-monotonic dependence on the growth rates[27]. We do not intend to explicitly discuss the growth rate-dependent manner of DnaA expression, as the growth rate is not experimentally altered in our work.

As mentioned above, the sequestration of oriC is a rather efficient mechanism preventing re-initiation, which determines the maximum number of replisomes and oriCs that can exist simultaneously on the same genome (see **Theorem 1.4**). The oriC and the *dnaA* regions become hemimethylated after replication initiation and stay hemimethylated for a considerable period where the closely spaced GATC sites in these regions are bound by SeqA proteins, which sequester oriC and the *dnaA* region[28,29]. During this period, DnaA cannot bind to oriC, and no *dnaA* is transcribed[28,29].

Furthermore, RIDA (regulatory inactivation of DnaA) is another predominant mechanism of negative regulation of DnaA function[30,31]. DnaA homolog protein Hda interacts with DnaA to promote hydrolysis of DnaA-bound ATP. This process requires a DNA-loaded clamp binding to Hda, which ensures RIDA is tightly coupled to active replication. *datA* loci can also mediate DnaA-ATP hydrolysis of DNA bound protein, a process named DDAH, whose contribution to DnaA-ATP hydrolysis is relatively small[32,33]. Interestingly, there are also two identified chromosomal DnaA-binding boxes that can catalyze DnaA-ADP rejuvenation (i.e., $\mathrm{DnaA_{ADP}} \rightarrow \mathrm{DnaA_{ATP}}$), known as DARS1 and DARS2 (DnaA-reactivating sequence 1 & 2)[15]. However, the cellular factors that regulate DARS activity are as yet unidentified.

ii. Mathematical description of DnaA regulations

In summary, five processes negatively regulate DnaA function: (1) DnaA negative autoregulation, (2) Binding of SeqA, (3) datA locus titration, (4) RIDA and (5) DDAH, and one positive regulation, DARS. A major challenge of modeling DNA replication initiation based upon these processes is limited quantitative knowledge about the kinetic parameters. We will later return to this and show how these processes could be merged in mathematical considerations to overcome the difficulty in parameter selection. For the regulation part of replication initiation modeling, we extend the DnaA-titration model first developed by Hansen et al.[34] with all the information above as major processes:

**a. Transcription of *dnaA* gene:**

$$\emptyset \rightarrow mRNA\ of\ dnaA \qquad Eq.\ 3.1$$

According to *Eq. 2.17*, the kinetic equation is

$$\frac{dm_{dnaA}}{dt} = \frac{v_{m,dnaA} g_{dnaA} \alpha_f R}{R + gK_t} \cdot \frac{K_{DnaA}^m}{K_{DnaA}^m + [DnaA_{ATP}]^m} - \frac{m_{dnaA}}{\tau_{m,dnaA}} \qquad Eq.\ 3.2$$

where the autorepression of DnaA-ATP in a cooperative manner $m > 1$ is involved. Meanwhile, transcription is completely shut off when *dnaA* region is sequestered by SeqA, which can be involved in calculating active gene copy number (see the binary tree part).

**b. Translation of DnaA**

$$mRNA\ of\ dnaA \rightarrow DnaA \qquad Eq.\ 3.3$$

whose kinetic equation is

$$\frac{dDnaA}{dt} = \sigma_{DnaA} v_e m_{DnaA} \frac{Rb - Rb'}{Rb - Rb' + mK_e} \qquad Eq.\ 3.4$$

where DnaA is a stable protein whose degradation rate is neglected.

**c. Binding of ATP or ADP to DnaA**

$$ATP + DnaA \overset{K_{ATP}}{\longleftrightarrow} DnaA_{ATP} \qquad Eq.\ 3.5$$

$$ADP + DnaA \overset{K_{ADP}}{\longleftrightarrow} DnaA_{ADP} \qquad Eq.\ 3.6$$

The processes have been believed to be rather fast, so we hypothesized fast equilibrium here. Given the fact that cellular $\gamma = \frac{[ATP]}{[ADP]}$ is approximately a constant, we have

$$\frac{[DnaA_{ATP}]}{[DnaA_{ADP}]} = \gamma K_{ATP} K_{ADP} \qquad Eq.\ 3.7$$

It is imaginable that a newly-synthesized DnaA protein immediately binds with either ADP or ATP. Therefore, joining *Eq. 3.7* with *Eq. 3.2* and *Eq. 3.4*, we have

$$\frac{dDnaA_{ATP}}{dt} = \sigma_{DnaA} v_e m_{DnaA} \frac{\gamma K_{ATP} K_{ADP}}{1 + \gamma K_{ATP} K_{ADP}} \frac{Rb - Rb'}{Rb - Rb' + mK_e} \qquad Eq.\ 3.8$$

$$\frac{dDnaA_{ADP}}{dt} = \sigma_{DnaA} v_e m_{DnaA} \frac{1}{1 + \gamma K_{ATP} K_{ADP}} \frac{Rb - Rb'}{Rb - Rb' + mK_e} \qquad Eq.\ 3.9$$

**d. Replicative inactivation of DnaA**

$$DnaA_{ATP} + RepFork \rightarrow DnaA_{ADP} \qquad Eq.\ 3.10$$

whose kinetic equation is written as

$$r_{RIDA} = k_{RIDA} [DnaA_{ATP}] n_{fork} \qquad Eq.\ 3.11$$

**e. DnaA binding and titration**

$$DnaA_{ATP} \rightarrow datA \qquad Eq.\ 3.12$$

$$DnaA_{ATP} \rightarrow \text{oriC high affinity boxes} \qquad Eq.\ 3.13$$

$$DnaA_{ATP} \rightarrow \text{oriC low affinity boxes} \qquad Eq.\ 3.14$$

The last two binding processes are believed to determine the initiation probability directly. Based upon these, we now enter the trigger part of DNA replication initiation, where we would discuss how to determine the initiation probability from the binding pattern in oriC.

iii. Two possible models for replication initiation

An intuitive hypothesis for DnaA accumulation at the oriC region to trigger initiation is that every DnaA box has to be occupied. This model has been suggested elsewhere to reproduce the phenomenological law of initiation mass. For convenience, we term this model DnaA random filling model (DRFM) because DnaA boxes are indistinguishable with respect to their roles in replication initiation. The initiation mass indicates that replication initiation occurs when cell mass per oriC reaches a certain threshold. The simplest understanding of this is that number of occupied DnaA boxes scales with the number of DnaA proteins in the cell. If intracellular DnaA concentration is approximately a constant, the initiation mass law can be readily implemented. However, the model obviously encounters challenges when we experimentally blocked different DnaA boxes. The order of replication inhibition strength is R1>R5M>R4>C3>I2. In DRFM, there is no difference among different boxes except for their binding affinities towards DnaA, which apparently are not the determinants for strength order (R1 and R4 are high-affinity boxes while R5M, C3, and I2 are all low-affinity boxes). Actually, with the hypothesis of DRFM, targeting high-affinity boxes should be less efficient than target low ones, as dCas9 is less competent towards high-affinity boxes. This is consistent with the fact the high-affinity boxes are occupied by DnaA most of the time. We simulated the DRFM *in silico* and found a pronounced weaker effect of high-affinity boxes (see below). This suggests that DnaA boxes should be further differentiated from each other from more perspectives.

It is noteworthy that DnaA proteins bound to DnaA boxes on different sides of the R2 box are in the opposite direction. On the left side of the R2 box, R1, R5M, τ2, I1, and I2 in the sequence are arranged rightward, while on the right side of the R2 box, C3, C2, I3, C1, and R4 are arranged leftward. Previous studies have identified that DnaA in the same direction can form oligomers in the oriC region, suggesting that there might be two oligomers forming independently[14]. From the mechanics perspective, DnaA oligomers provide double-stranded DNA with a torsion, which is positively related to the length of oligomers. Given that, we propose a double DnaA oligomer model (DDOM), where two DnaA monomers first occupy R1 and R4 boxes to nucleate two DnaA oligomers, and two DnaA oligomers jointly contribute to DNA unwinding at the DNA unwinding element (DUE). The contribution of an oligomer is supposed to be positively correlated with its length, and two oligomers contribute differently to DNA unwinding. Qualitatively, DDOM is sufficient to explain the observation. In the left oligomer, because I2 is behind R5M, which is further behind R1, the inhibitory effects from strong to weak are R1, R5M and I2, and the same is true for R4 and C3 in the right oligomer. Moreover, the fact that R1 has a more substantial effect than R4 indicates a larger impact on replication initiation for a replication origin without the left oligomer, which is consistent with previous work that deleted the left and right parts of oriC[35,36].

Based on these analyses, we further give a clear description of DnaA-driven replication initiation in DDOM. In our experiments, we do not perturb the factors other than DnaA binding, and thereby we assume that DnaA-ATP occupancy is the only determinant of the timing of initiation. Given that, we first define an occupancy vector to describe if a certain box is occupied or not:

> **Definition 3.1** (The occupancy vector). Given a replication origin with $\boldsymbol{N}$ DnaA binding boxes, for $\boldsymbol{i} = \mathbf{1}, \mathbf{2}, \mathbf{3}, \cdots, \boldsymbol{N}$, define $\boldsymbol{\omega_i} = \mathbf{1}$ if the $\boldsymbol{i}^{\text{th}}$ box is occupied by DnaA and $\boldsymbol{\omega_i} = \mathbf{0}$ if not. At the moment $\boldsymbol{t}$, the occupancy vector of the replication origin is defined as $\boldsymbol{\Omega}(\boldsymbol{t}) = [\boldsymbol{\omega_1}(\boldsymbol{t}), \boldsymbol{\omega_2}(\boldsymbol{t}), \cdots, \boldsymbol{\omega_N}(\boldsymbol{t})]$.

For a fixed occupancy vector, the probability of DNA replication initiation is supposed to be determined. Considering that the process can be treated as a chemical reaction $\text{DNA} \rightarrow \text{unwound DnaA}$, we use a definition of Poisson-type stochastic process for DNA replication initiation:

> **Assumption 3.1** (DNA replication initiation). For a given replication origin, define a random variable $\boldsymbol{I} = \mathbf{1}$ if DNA replication initiation occurs and $\boldsymbol{I} = \mathbf{0}$ if not. Given $\boldsymbol{\kappa}$, a variable that only depends on occupancy vector $\boldsymbol{\Omega_N}$ of N DnaA boxes by DnaA proteins, i.e., $\boldsymbol{\kappa} \equiv \boldsymbol{\kappa}(\boldsymbol{\Omega_N})$, if $\boldsymbol{\tau} \in [\mathbf{0}, \boldsymbol{\tau_m}]$ is the observation time during which $\boldsymbol{\Omega_N}$ remains unchanged, DNA replication initiation is a stochastic process with an intensity $\boldsymbol{\kappa}$ which follows: (1) $\boldsymbol{I}(\boldsymbol{\tau} = \mathbf{0}) = \mathbf{0}$; (2) For $\forall \boldsymbol{t} \geq \mathbf{0}$, given $\boldsymbol{h} \rightarrow \mathbf{0}^{+}$, $\boldsymbol{P}(\boldsymbol{I}(\boldsymbol{h}) = \mathbf{1}) = \boldsymbol{\kappa h} + \boldsymbol{o}(\boldsymbol{h})$; (3) The process is an independent and stationary increment process.

**Assumption 3.1** is the foundation for numerical simulations of replication initiation. Actually, with a period $t$ divided into $n$ equal intervals, when $n \rightarrow \infty$, we have

$$P(I(t)) = P\left(I\left(\frac{t}{n}\right) = 0\right)^n = \left[1 - P\left(I\left(\frac{t}{n}\right) = 1\right)\right]^n \rightarrow e^{-\kappa t} \qquad Eq.\ 3.15$$

suggesting that the waiting time follows an exponential distribution with mean $\frac{1}{\kappa}$. Therefore, $\kappa$ is the rate of replication initiation. Typically, if a DnaA box $i$ is occupied by dCas9 and the permitted occupancy of DnaA-ATP at most is $\Omega_{N,i}$, given that the dissociation rate of dCas9 is $k_{-}$, the timing of initiation will be dominated by dCas9 dissociation if $\kappa(\boldsymbol{\Omega}_{N,i}) \gg k_{-}$, or by the probability of DNA unwinding with dCas9 still occupying the oriC if $\kappa(\boldsymbol{\Omega}_{N,i}) \ll k_{-}$.

Through the transition state theory, the reaction rate of replication initiation $\lambda$ is supposed to follow

$$\kappa = Ae^{\frac{E_0 - E^{\ddagger}}{kT}} \qquad Eq.\ 3.16$$

It is understandable that torsion of DnaA oligomers increases the instability of DNA and thus facilitates its unwinding. Adding monomers to the oligomer thus increases $E_0$. Based on that, by integrating the constants, we define initiation energy which directly correlates with the number of monomers:

**Assumption 3.2** (Initiation energy). Given a DNA replication initiation process with an intensity $\boldsymbol{\kappa}$, define the initiation energy as $\boldsymbol{E_{ori} = \ln \kappa}$. $\boldsymbol{E_{ori}}$ has a linear relation with the occupancy vector $\boldsymbol{\Omega_N}$ of the replication origin by $\boldsymbol{E_{ori}(t) = \Omega_N(t) K_{ori} + K_0}$, where $\boldsymbol{K_{ori}}$ is a time-independent column vector with $\boldsymbol{N}$ non-negative elements and $\boldsymbol{K_0}$ is a constant number.

Physically, $\boldsymbol{K_{ori}}$ refers to the contribution of each box to DNA unwinding. Although the linear relation by **Assumption 3.2** is a rough description of the replication initiation complex, it provides a good insight on the physical model of DnaA-driven DNA replication initiation and order of contributions of different boxes. In the end, we introduce DDOM:

**Assumption 3.3** (DDOM). $\boldsymbol{N}$ DnaA boxes in the replication origin region are composed of two continuous sequences with $\boldsymbol{p}$ and $\boldsymbol{q}$ boxes respectively

$(\boldsymbol{p} + \boldsymbol{q} = \boldsymbol{N})$, i.e., $\boldsymbol{\Omega_N(t)} = [\boldsymbol{\Omega_p(t)}, \boldsymbol{\Omega_q(t)}]$. For $\forall \boldsymbol{t} \in \mathbb{R}$, in either $\boldsymbol{\Omega_p(t)}$ or $\boldsymbol{\Omega_q(t)}$, with $\boldsymbol{i} = \boldsymbol{2}, \boldsymbol{3}, \cdots, \boldsymbol{p}$ ($\boldsymbol{q}$ for $\boldsymbol{\Omega_q(t)}$), if $\boldsymbol{\omega_i(t)} = \boldsymbol{1}$, there must be $\boldsymbol{\omega_{i-1}(t)} = \boldsymbol{1}$.

This requirement is the only difference between DDOM and DRFM. In other words, DDOM requires that a DnaA box can be occupied only when the box before it has already been occupied. Maybe a more accurate expression is that a box can be bound by DnaA monomer at any time, but it does not contribute to DNA unwinding until its adjacent box is also bound. For *E. coli*, the corresponding boxes of the occupancy vector can be specified as $\boldsymbol{\Omega_p(t)}: [R1, R5M, \tau 2, I1, I2]$ and $\boldsymbol{\Omega_q(t)}: [R4, C1, I3, C2, C3]$. It is noteworthy that the R2 box is not involved here, for which there are mainly two reasons. First, the R2 box was believed to rescue normal DNA unwinding when DnaA boxes on its left side are entirely deleted, suggesting that R2 might be an alternative site for nucleating an oligomer, though apparently in a much less efficient way[36]. Considering the process is not yet clear, and it has been reported that the high-affinity boxes are occupied most of the time[37,38], it is proper to treat the occupied state of the R2 box as an invariant in this model. Secondly, the R2 box is not a target for *CRISPRori* in our experiments, and thereby neglecting the R2 box will not affect the simulation.

**Supplementary Note 4. *CRISPRori* inhibition on the DnaA boxes**

i. Four-state binding model

To elaborate on the dose-dependent inhibitory effect by *CRISPRori*, we estimated the time delay of replication initiation by a four-state competitive binding model between dCas9 and DnaA to the *oriC* sites.

$$BD \overset{k_{\pm1}}{\longleftrightarrow} B \overset{k_{\pm2}}{\longleftrightarrow} BA \longrightarrow Initiation \qquad Eq.\ 4.1$$

where $D$ denotes dCas9, $B$ denotes DnaA box, and $A$ denotes DnaA. The formula is able to cover both two possible pathways for an initiation event to occur: $B \rightarrow BA$ and $B \rightarrow BD \rightarrow B \rightarrow BA$. To calculate how long a dCas9 protein delays an initiation event, we treated *Eq. 4.1* as a one-dimensional random walk and calculated the average time taken from $BD$ to $BA$. For mathematical tractability, we apply the balanced growth assumption, which approximates the concentration of $D$ and $A$ as constants.

In the one-dimensional random walk, the waiting time of state $i$ should be an exponential distribution with a mean value of $\frac{1}{\sum_j k_{ij}}$ where $k_{ij}$ is the observatory rate constant of reaction $i \rightarrow j$. According to the strong Markov property of the continuous-time Markov chain, we have:

$$\begin{cases} \langle T_{BD\to BA} \rangle = \dfrac{1}{k_1} + \langle T_{B\to BA} \rangle \\ \langle T_{B\to BA} \rangle = \dfrac{1}{k_{-1}D + k_2 A} + \dfrac{k_{-1}D}{k_{-1}D + k_2 A} \langle T_{BD\to BA} \rangle \end{cases} \qquad Eq.\ 4.2$$

Solve *Eq. 4.2* and we obtain the mean delay of an initiation event upon dCas9 binding:

$$\langle T_{BD\to BA} \rangle = \frac{k_{-1}}{k_1 k_2} \frac{D}{A} + \frac{1}{k_2 A} + \frac{1}{k_1} \qquad Eq.\ 4.3$$

Without considering the binding order of DnaA to oriC region as demonstrated in the preceding section, we can dissect the initiation probability into several events highlighting the impact of dCas9 occupancy:

$E_1$: DnaA replaces dCas9;

$E_2$: All boxes except for the target box are occupied by DnaA;

$E_3$: dCas9 binds the target box;

$E_4$: All boxes are occupied by DnaA.

Hence, the initiation probability should be:

$$P_{init} \propto P(E_4) = P(C)P(AB|C) + P(\overline{C})P(D|\overline{C}) \qquad \textit{Eq. 4.4}$$

Substituting *Eq. 4.3* into *Eq. 4.4*, we have

$$\langle T_{init} \rangle = P(C)\langle T_{BD\to BA} \rangle + P(\overline{C}) \cdot 0$$

$$= \langle T_{BD\to BA} \rangle \int_0^{\Delta\tau_i} k_{-1} D e^{-k_{-1}Dt} \mathrm{d}t$$

$$= (1 - e^{-k_{-1}D\Delta\tau_i}) \left( \frac{k_{-1}}{k_1 k_2} \frac{D}{A} + \frac{1}{k_2 A} + \frac{1}{k_1} \right) \qquad \textit{Eq. 4.5}$$

where $\Delta\tau_i$ is the time interval between two successive initiation events.

ii. Stochastic simulation

Based on the discussions above, we further performed a simple stochastic simulation using the Gillespie algorithm to evaluate the heterogeneity of cell responses to *CRISPRori* treatment. For each simulation, a $B \to BD$ reaction is firstly simulated to determine if a dCas9 protein can bind to oriC within $\Delta\tau_i$. Expression curve of dCas9 induced by arabinose is experimental measured and fitted to Hill function:

$$D = \frac{\beta_D \left(\frac{c_D}{K_D}\right)^n}{1 + \left(\frac{c_D}{K_D}\right)^n} \qquad \textit{Eq. 4.6}$$

Theoretical predictions showed the average initiation delay time increased with the inducer concentration until it plateaued at ~200min (at ~0.3% arabinose), which was close to the dissociation time of dCas9 **(Supplementary Fig. 5C)**[39]. This is because the dissociation term $1/k_1$ is much longer than the other two in *Eq. 4.3*:

$$\langle T_{init} \rangle \approx \frac{1}{k_1}(1 - e^{-k_{-1}D\Delta\tau_i}) \xrightarrow{D\to+\infty} \frac{1}{k_1} \qquad \textit{Eq. 4.7}$$

During this process, the population underwent a transition from a bimodal phase to a homogenous phase in terms of single-cellular division time, which was explained by the diffusion and dissociation of dCas9, respectively. At low inducer concentrations, the diffusion of dCas9 is the rate-limiting step. The probability that dCas9 preceded DnaA to bind *oriC* is low, generating two sub-populations—cells with or without dCas9 bound to *oriC* **(Supplementary Fig. 5A,B)**. Experimentally, populations were partitioned into normal-sized and filamentous cells in groups of low inducer concentrations or in the early inducing stage **(Supplementary Fig. 5D)**. As the inducer concentration or induction time increased, filamentous cells became the majority, and heterogeneity in cell length disappeared since the rate of dCas9 dissociation from *oriC* sites began to govern the inhibition period once dCas9 loaded onto *oriC*.

### Numerical simulation protocol

#### Supplementary Note 5. Algorithm

A major challenge in bridging microcosmic molecular mechanisms with phenomenological observations is to simulate gene expression and DNA replication jointly. To address this issue in numerical simulation, an *E. coli* cell was divided into two primary systems: the reaction and genome systems. Two corresponding encapsulated objects were created for each, where information exchange between them was highly required to describe the impact of DNA replication on gene expression and vice versa.

##### i. Langevin equations

We denote the reaction system, which is comprised of a set of Langevin equations by $\mathbb{L}$. $\mathbb{L}$ involves transcription, translation, mRNA degradation, protein degradation of RNAP, ribosomes, DnaA-ATP/ADP, FtsZ, and plasmid-borne GFP. For simplicity, we considered the plasmid copy number control as a steady instead of a dynamical process by assuming that plasmid copy number scaled with the total number of proteins. For proteins encoded by several genes, like RNAP and ribosome, only one gene was selected to represent the whole protein (see below). Besides, datA titration, binding of oriC by DnaA or dCas9, dissociation of dCas9, and replicative inactivation of DnaA (RIDA) are all simulated by the progressive stepwise solution of equations as mentioned before (see the general theoretical framework section) together with a Langevin Gaussian noise term.

##### ii. Bacterial genome

We denote the bacterial genome system by $\mathbb{G}$. Functions in the genome system could be divided into two classes: genome-associated events and functions of random binding of dCas9 and DnaA. The genome, as mentioned before, is described by a binary tree where genome-associated events, like DNA replication initiation and termination, are direction alterations of the tree structure (see **The data structure of a bacterial**

**genome**). As for random binding of dCas9 and DnaA whose reaction extents are determined by the Langevin equations in $\mathbb{L}$, the $\mathbb{G}$ system performed a uniform random sampling among all available boxes on all genomes to specify which box to be bound. Clarification of specific boxes to bind in each loop of the update is required for calculating the DNA replication initiation probability as discussed in **Two possible models for replication initiation**.

iii. The criterion for cell division

In this model, we adopt the most straightforward hypothesis to judging cell division that has been developed and experimentally validated to explain the observation of "adder"[40]. FtsZ, the protein that forms the division ring of *E. coli*, was assumed to accumulate to a certain threshold to trigger cell division. To ensure the genome integrity in offspring cells, bacteria employed a nucleoid occlusion mechanism to prevent cell division when only one replicating nucleoid exists in cell[41,42]. Hence, the criterion for cell division should be

$$FtsZ > threshold \ \textbf{and} \ \# \ Nucleoids \geq 2 \qquad Eq.\ 5.1$$

Upon division, cell volume undergoes asymmetric division following the binary random distribution. For a cell with $N = 2k + 1\ (k > 1)$ nucleoids, $k$ and $k + 1$ nucleoids were allocated toward the smaller and bigger cells, respectively.

iv. Coordination of gene expression, DNA replication, and cell division

To implement progressive stepwise simulation, variables had to be passed to each other between $\mathbb{L}$ and $\mathbb{G}$. In **Table 5.1**, we summarize the functions to exert in black, which require variables in blue.

**Table 5.1.** Functions that require variables from another system

| $\mathbb{L}$ | $\mathbb{G}$ |
|---|---|
| Transcription | Gene copy number, total gene copy number, and sequestration state |
| RIDA | Number of replication forks |

| | |
|---|---|
| dCas9 binding | Number of available target boxes and sequestration state |
| DnaA binding | Number of available DnaA boxes and sequestration state |
| DnaA titration | Number of available datA boxes |
| dCas9 dissociation | Number of target boxes blocked |
| Number of DnaA bound | DnaA boxes occupancy & judge replication initiation |
| Number of dCas9 bound | Target box blocking |

In every loop of the simulation, the process is as follows:

(i) Update all reactants in $\mathbb{L}$ in $\Delta t$;

(ii) Return the number of high-affinity boxes bound $N_h$, low-affinity boxes bound $N_l$, dCas9 binding $N_{\mathrm{on}}$, dCas9 dissociated $N_{\mathrm{off}}$, and datA binding $N_t$;

(iii) Uniformly distribute $N_h, N_l, N_{\mathrm{on}}, N_{\mathrm{off}}, N_t$ to all genomes $\{\mathbb{G}_{\mathbf{1}}, \mathbb{G}_{\mathbf{2}}, \cdots, \mathbb{G}_{\mathbf{n}}\}$ based on the number of available or blocked boxes;

(iv) For certain genome $\mathbb{G}_{\mathbf{i}}$ in $\{\mathbb{G}_{\mathbf{1}}, \mathbb{G}_{\mathbf{2}}, \cdots, \mathbb{G}_{\mathbf{n}}\}$:

a) Perform progressive binding of a certain type of box $b$ with the total number $N_{b,i}$. For every DnaA binding to oriC (i.e., either high-affinity or low-affinity box), calculate the initiation probability and judge if the initiation occurs. If it does, release all DnaA and dCas9 in the oriC region and switch to sequestration state.

b) Every replisome moves a step forward. If particular pair of replisomes arrives at the terminus, divide $\mathbb{G}_{\mathbf{i}}$ into two and update all genomes to $\{\mathbb{G}_{\mathbf{1}}, \mathbb{G}_{\mathbf{2}}, \cdots, \mathbb{G}_{\mathbf{n+1}}\}$.

(v) If the number of intracellular FtsZ reaches threshold and $\#\{\mathbb{G}\} \geq 2$, distribute all reactants and $\{\mathbb{G}\}$ into offspring cells. In practice, we only retain one of the two offspring cells to track the lineage.

#### Supplementary Note 6. Parameters

**Table 6.1.** Table of parameters used in the model.

| Parameter | Physical meaning | Value | Unit |
|---|---|---|---|
| $\alpha_{inact}$ | Fraction of inactive ribosomes in total proteins: $\alpha_{inact} = R'/P$, see *Eq. 2.10.* | 0.002[a] | 1 |
| $\alpha_f$ | Fraction of free RNAP in total RNAP. See *Eq. 2.2.* | 0.22[1] | 1 |
| $K_{ATP}$ | Equilibrium constant of DnaA binding with ATP. See *Eq. 3.5.* | 0.05[b] | 1 |
| $K_{ADP}$ | Equilibrium constant of DnaA binding with ADP. See *Eq. 3.6.* | 0.0125[b] | 1 |
| $v_e$ | Ribosome elongation rate. See *Eq. 2.6* | 2.88[c] | $\text{min}^{-1}$ |
| $\lambda$ | Exponential growth rate | 0.0170[d] | $\text{min}^{-1}$ |
| $K_e$ | Number of ribosomes half-saturating an mRNA. See *Eq. 2.7.* | 7.7[e] | 1 |
| $K_t$ | Number of RNAP half-saturating the transcription. See *Eq. 2.1.* | 0.4[f] | 1 |
| $k_{hbox}$ | DnaA-ATP binding rate on high-affinity boxes | 0.06[43] | $\text{min}^{-1}$ |
| $k_{lbox}$ | DnaA-ATP binding rate on low-affinity boxes | 0.006[43] | $\text{min}^{-1}$ |
| $k_{off}$ | dCas9 dissociation rate | 0.005[39] | $\text{min}^{-1}$ |
| $k_{on}$ | dCas9 binding rate | 0.01[39] | $\text{min}^{-1}$ |
| $k_{RIDA}$ | Rate of RIDA. See *Eq. 3.11.* | 0.2[g] | $\text{min}^{-1}$ |
| $m$ | Hill coefficient of DnaA autorepression. See *Eq. 3.2.* | 3[h] | 1 |
| $K_{DnaA}$ | Number of DnaA that reduces its transcription to half | 1000[h] | 1 |
| $v_{DnaA}$ | Maximal transcription rate of DnaA. | 3[i] | $\text{min}^{-1}$ |
| $v_{FtsZ}$ | Maximal transcription rate of FtsZ. | 5[i] | $\text{min}^{-1}$ |

| | | | |
|---|---|---|---|
| $\nu_{GFP}$ | Maximal transcription rate of plasmid GFP. | 10[*] | $\text{min}^{-1}$ |
| $\nu_R$ | Maximal transcription rate of RNAP. | 3[i] | $\text{min}^{-1}$ |
| $\nu_{Rb}$ | Maximal transcription rate of DnaA. | 30[1,i] | $\text{min}^{-1}$ |
| $\nu_T$ | Maximal transcription rate of all proteins on average. | 1[i] | $\text{min}^{-1}$ |
| $\gamma$ | Ratio of intracellular ATP to ADP. See *Eq. 3.7.* | 3.2[44] | 1 |
| $\sigma_{DnaA}$ | Maximal ribosomal density on the mRNA of DnaA. See *Eq. 2.8.* | 3[j] | $\text{Knt}^{-1}$ |
| $\sigma_T$ | Maximal ribosomal density of all proteins on average. See *Eq. 2.8.* | 10[45] | $\text{Knt}^{-1}$ |
| $t_{sequest}$ | Sequestration time after an initiation event. | 10[13] | min |
| $\tau_{m,DnaA}$ | mRNA degradation time of DnaA. | 13.6[46] | min |
| $\tau_{m,FtsZ}$ | mRNA degradation time of FtsZ. | 16.3[47] | min |
| $\tau_{m,R}$ | mRNA degradation time of RNAP. | 9.8[†] | min |
| $\tau_{m,Rb}$ | mRNA degradation time of ribosomes. | 17.5[46] | min |
| $\tau_{m,GFP}$ | mRNA degradation time of GFP. | 9.8 | min |
| $\tau_{m,T}$ | mRNA degradation time of all proteins on average. | 9.8[46] | min |
| $\tau_{p,FtsZ}$ | Degradation time of FtsZ. | 105[48] | min |
| $A_0$ | Initial DnaA number. | 1000[49] | 1 |
| $Z_0$ | Initial FtsZ number. | 10000[50] | 1 |
| $Z_t$ | FtsZ number threshold triggering division. | 14000[k] | 1 |
| $R_0$ | Initial RNAP number. | 8400[1] | 1 |
| $Rb_0$ | Initial ribosome number. | 67500[51] | 1 |
| $M_0$ | Initial mRNA number. | 8400[52] | 1 |
| $P_0$ | Initial protein number. | 7500000[53] | 1 |
| $C$ | *C*-period. | 40[54] | min |

[*] This is an artificial number because GFP is plasmid-borne, where the promoter can be flexibly altered.

[†] RNAP and GFP mRNA degradation times directly take the value of average number as they are not measured by the previous work.

| | | | |
|---|---|---|---|
| $G$ | Total number of genes. | 4500 | 1 |
| $N_t$ | DnaA boxes in datA | 200[13] | 1 |
| $L_{DnaA}$ | Genomic location of *dnaA*. | 0.0185[‡] | 1 |
| $L_{FtsZ}$ | Genomic location of *ftsZ*. | 0.354 | 1 |
| $L_R$ | Genomic location of *rpoC*, one of genes encoding RNAP. | 0.1127 | 1 |
| $L_{Rb}$ | Genomic location of *rpsA*, one of genes encoding the ribosome. | 0.7233 | 1 |
| $L_{datA}$ | Genomic location of *datA* | 0.201 | 1 |

**Notes:**

a. Molecular fraction of inactive ribosomes $\alpha_{inact}$.

It has been reported that $r = r_0 + \lambda \cdot \text{const.}$, where $r$ denotes the mass ratio of RNA to protein and $r_0 \approx 0.1$ (see ref.[55]). Given the fact that (1) 86% of total RNA is rRNA in *E. coli*[54], (2) total mass of three rRNA molecules is $1.54 \times 10^6$Da, and (3) average protein length is 325 amino acids[56], we have

$$\alpha_{inact} = \frac{0.1 \times 0.86 \times 325 \times 110^{\S}}{1.54 \times 10^6} \approx 0.002 \qquad \textit{Eq. 6.1}$$

b. Equilibrium constants of DnaA binding ATP and ADP $K_{ATP}$ and $K_{ADP}$.

Previous works have measured that $\mathrm{K_D}$ for ATP and ADP are 10 to 30 nM and 60 to 100 nM, respectively[15]. Given an average cell volume of $2\ \mu\mathrm{m}^3$ and take the middle number 20 nM and 80 nM, we have

$$K_{ATP} = \frac{1}{20 \times 10^{-9} \times 2 \times 10^{-15} \times 6.02 \times 10^{23}} \approx 0.05 \qquad \textit{Eq. 6.2}$$

$$K_{ADP} = \frac{K_{ATP}}{4} \approx 0.0125 \qquad \textit{Eq. 6.3}$$

c. Ribosome elongation rate $v_e$

[‡] All gene locations are from NCBI.

[§] Average molecular weight of an amino acid is about 110 Da.

The typical translation rate of a ribosome is 15 amino acids/sec, and recalling that the average protein length is 325 amino acids[56], we have

$$v_e = \frac{15 \times 60}{325} \approx 2.8\ \text{protein/min} \qquad \textit{Eq. 6.4}$$

d. Exponential growth rate $\lambda$

$\lambda$ is linearly fitted to the lineage cell length data of the negative control group from the measurement of microfluidic systems. Namely, the doubling time is $\tau = \frac{\ln 2}{\lambda} \approx 40\ \text{min}$.

e. Number of ribosomes half-saturating an mRNA $K_e$

Recalling that our derivation of *Eq. 2.28* is consistent with previously reported $r = r_0 + \lambda \cdot \mathrm{c}$, $K_e$ can be estimated from the previous measurement that $\mathrm{c} \approx 0.23$ hrs[55]. Again, the mass ratio $r$ and the molecular ratio $\Phi_{Rb}$ should have the following relation:

$$\Phi_{Rb} = r \cdot \frac{0.86 \times 325 \times 110}{1.54 \times 10^6} \approx 0.02r \qquad \textit{Eq. 6.5}$$

Hence,

$$\frac{K_e}{v_e \langle \sigma \rangle} \approx 0.02\mathrm{c} \approx 0.276\ \text{min} \qquad \textit{Eq. 6.6}$$

With $\langle \sigma \rangle \approx 10$ and $v_e \approx 2.8\ \text{min}^{-1}$, we have:

$$K_e = 0.276 \times 10 \times 2.8 \approx 7.7 \qquad \textit{Eq. 6.7}$$

f. Number of RNAP half-saturating transcription $K_t$

Previous work has shown that an E. coli cell with a growth rate of $1.5\ \text{doublings/h}$ has the time interval between two transcription firing of *Pspc* promoter as twice long as the minimum time interval[1]. This growth rate is close to ours since the doubling time of cells in our experiments is 40 min. Therefore, it should be reasonable to calculate the average active RNAP per promoter directly:

$$K_t \approx \frac{\#\ Active\ RNAP}{\#\ Promoters} = \frac{1848}{4500} \approx 0.4 \qquad \textit{Eq. 6.8}$$

g. RIDA rate $k_{RIDA}$

There are about 1,000 DnaA molecules in an *E. coli* cell. All DnaA-ATP molecules have to be inactivated at least once to "reset" the replication state for every replication cycle. Moreover, every oriC and datA combination can accommodate about 210 DnaA-ATP molecules at most, and we assume that this number reduces by half when averaging over the replication cycle. Hence, the RIDA rate can be roughly estimated as

$$k_{RIDA} \approx \frac{1000}{\frac{210}{2} \times 40} \approx 0.2\ \mathrm{min}^{-1} \qquad \textit{Eq. 6.9}$$

h. Hill coefficient and half-saturating number in DnaA autorepression function, $m$ and $K_{DnaA}$

Previous in vitro study showed that three DnaA boxes in the dnaA promoter region preferentially bind DnaA-ATP[15], and thereby $m = 3$ is adopted. The real number could be greater if other boxes non-selectively binding DnaA-ATP and DnaA-ADP have contributions to the cooperative binding. There is lack of literature reporting $K_{DnaA}$, and considering the moderate effect of DnaA autorepression, we make a rather parsimonious assumption that $K_{DnaA}$ is comparable to the total number of DnaA in a normal-sized cell. Simulations show that $K_{DnaA}$ is not a sensitive parameter, and $\pm 25\%$ derivation from this value does not have remarkable impact on the results. However, complete removal of DnaA autorepression can influence the normal initiation mass (**Supplementary Fig. 14C,D**).

i. Maximal transcription firing rate of a certain gene $i$, $\nu_i$

In the analysis of balanced exponential growth, without consideration of gene transcription regulation, we show that

$$\phi_i^* = \frac{\sigma_i \mu_i^*}{\langle \sigma \rangle} = \frac{\sigma_i}{\langle \sigma \rangle} \frac{\nu_{m,i} \overline{\psi}_i}{\sum_j \nu_{m,j} \overline{\psi}_j} \propto \sigma_i \nu_{m,i} \qquad \textit{Eq. 6.10}$$

suggesting that given a fixed cell volume, the intracellular number of certain proteins should be proportional to the product of maximal ribosome density and its transcription firing rate. Since the transcription rate of ribosome protein promoter *Pspc* has been measured, the transcription rate of other genes can be estimated by

$$v_{m,i} \approx \frac{\#\,Protein\ i}{\#\,Ribosomes} \cdot \frac{\sigma_{Rb}}{\sigma_i} \cdot v_{Rb} \qquad Eq.\ 6.11$$

j. Maximal ribosome density on the mRNA of DnaA $\sigma_{DnaA}$

As mentioned before, the GUG start codon has about 3-fold lower translation initiation efficiency than AUG codon. Hence, the ribosome density is estimated to be $10/3 \approx 3$ ribosomes/Knt.

k. FtsZ threshold to trigger cell division, $Z_t$

As a matter of fact, the initial cell volume of simulation is set to be $3\ \mu m^3$, and to fit the division volume in NC group that is about $4.2\ \mu m^3$, initial FtsZ number is multiplied by 1.4 to ensure the division volume is close to the experiment.

### Discussions

#### Supplementary Note 7. Convergence rate of post-inhibition cell length by “adder” and “timer” laws

Let $L_b$ be the cell length at the start of a division cycle (i.e., birth length). For an *E. coli* lineage, define a series of $L_b$ of generations in sequence:

$$\{L_{b,0}, L_{b,1}, L_{b,2}, \cdots\} \qquad \textit{Eq. 7.1}$$

Mechanistically, if cell division is governed by FtsZ accumulation, cell size should follow the adder principle, which gives

$$L_{b,k} - \Delta L_{adder} = \left(\frac{1}{2}\right)^k (L_{b,0} - \Delta L_{adder}) \qquad \textit{Eq. 7.2}$$

$$\therefore \lim_{k\to\infty} L_{b,k} = \Delta L_{adder} \qquad \textit{Eq. 7.3}$$

The convergence rate of the $L_b$ should be

$$\lim_{k\to\infty} \frac{\left|L_{b,k+1} - \Delta L_{adder}\right|}{\left|L_{b,k} - \Delta L_{adder}\right|} = \frac{1}{2} \qquad \textit{Eq. 7.4}$$

Under exponential growth, time for a cell to finish an adder division cycle should be:

$$\Delta t_{adder} = \frac{1}{\lambda}\ln\left(1 + \frac{\Delta L_{adder}}{L_b}\right) \qquad \textit{Eq. 7.5}$$

However, for a cell where there is excessive FtsZ but only one nucleoid, its cell division is governed by the timing of replication termination. As a simplest situation, with a constant inter-initiation period of the replication cycle, cell size should follow a timer:

$$L_{b,k+1} = L_{b,k} \cdot \frac{e^{\lambda \Delta t_{init}}}{2} \qquad \textit{Eq. 7.6}$$

As long as the inter-initiation period satisfies that $\Delta t_{init} < \frac{ln\,2}{\lambda}$ = Doubling time, $L_b$ can converge to zero with a rate $\frac{e^{\lambda \Delta t_{init}}}{2} > \frac{1}{2}$. But undoubtedly, $L_b$ can never converge to zero because the amount of FtsZ can fall under the division threshold when cell volume is small enough. With a rough assumption that FtsZ concentration is a constant, *Eq. 7.6* holds only for a one-nucleoid cell whose $L_b > 2\Delta L_{adder}$. Assuming that that is an elongated cell with $L_{b,0} = n\Delta L_{adder} (n > 2)$, to make $L_b < 2\Delta L_{adder}$

through $k$ successive time division cycles, $k$ should follow

$$k > \frac{\ln \frac{n}{2}}{\ln \frac{2}{e^{\lambda \Delta t_{timer}}}} \qquad \textit{Eq. 7.7}$$

Through adder division cycles, $k$ should follow

$$k > \frac{\ln n - 1}{\ln 2} \qquad \textit{Eq. 7.8}$$

### Supplementary Figures

#### Supplementary Figure 1.

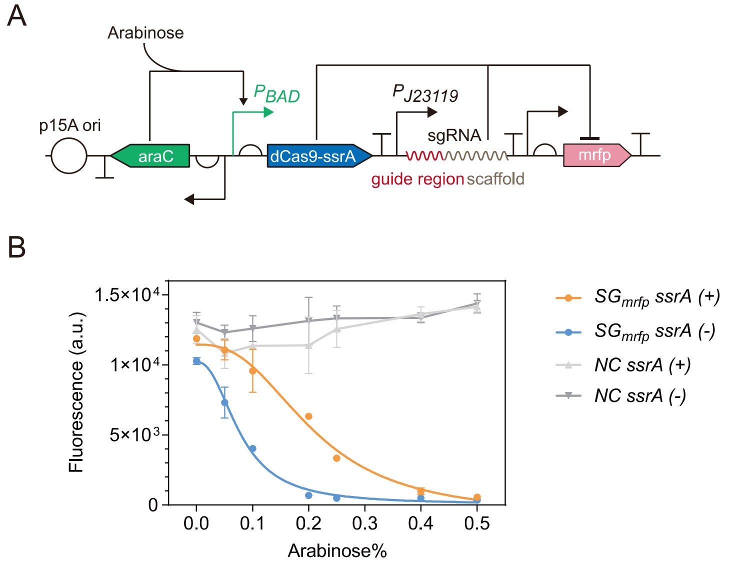

**Supplementary Figure 1. CRISPR-interference experiments with mRFP as the reporter.** Experimental results suggested the addition of a ssrA degradation tag to dCas9 yielded a more moderate inhibitory effect. (A) Plasmid construction of CRISPRi test. (B) NC: negative control. Error bars show s.d. of three experimental replicates.

**Supplementary Figure 2.**

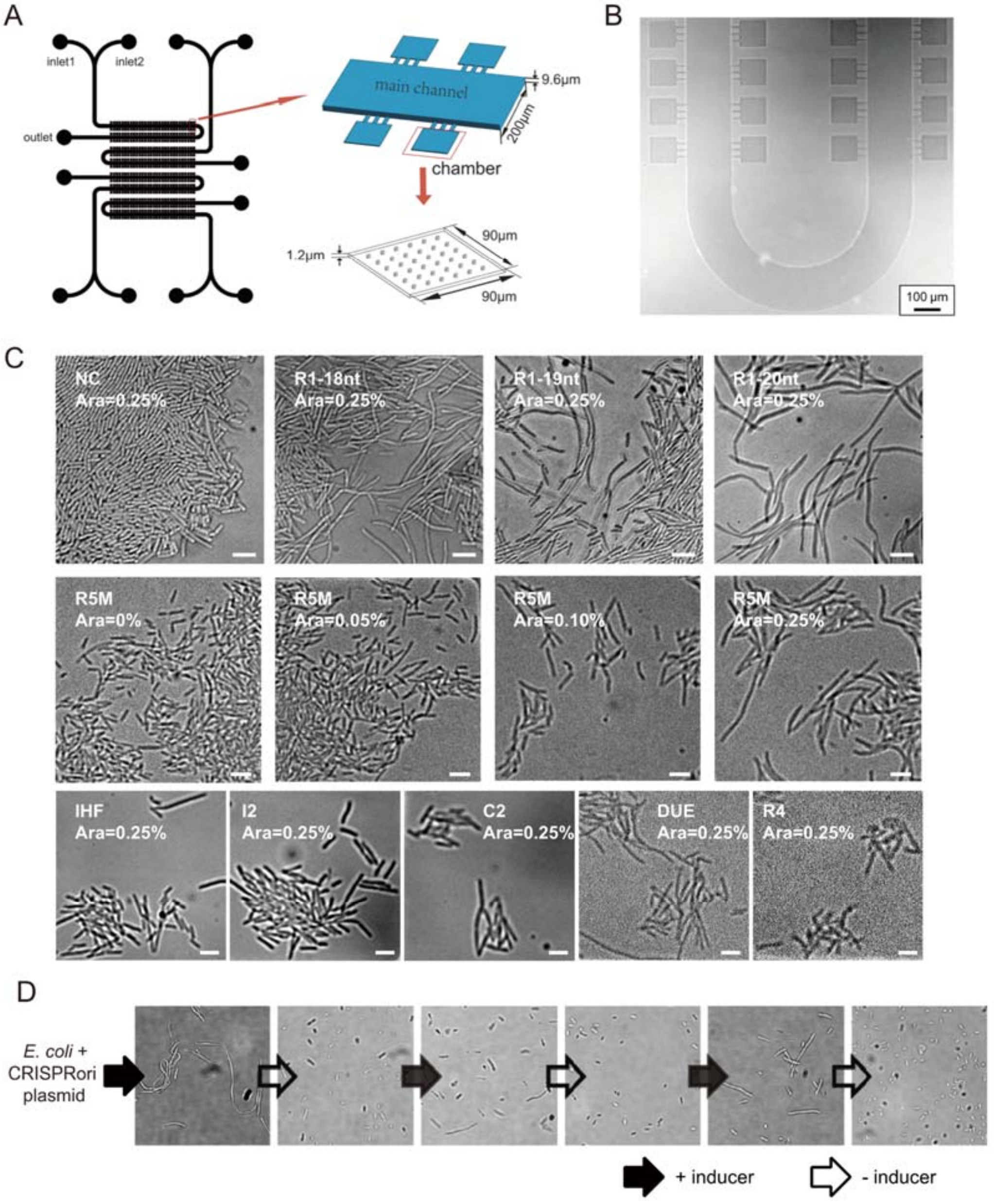

**Supplementary Figure 2. Experimental observations of cell morphologies in *CRISPRori* perturbed *E. coli* populations.** (A) Schematic of microfluidic device. (B) Microscopic photo of chambers in the microfluidic chip. (C) Time-lapse microscopic shots of growing *E. coli* cells in microfluidic chambers of different experimental groups. Scale bar = 4 μm. (D) Reversible initiation control. Cells were able to restore to normal morphologies in between episodes of *CRISPRori* induction.

**Supplementary Figure 3.**

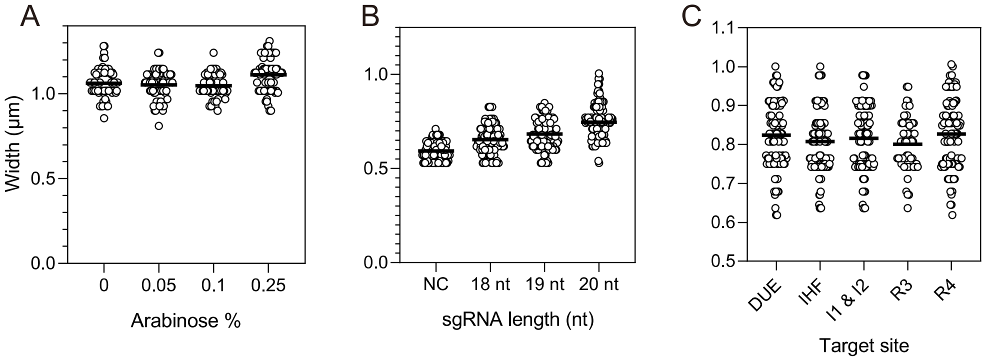

**Supplementary Figure 3. Cell widths were not significantly affected by *CRISPRori* expression.** In (B), the increase of mean cell widths was significant but slight compared to the dramatic changes observed in experimental manipulations of genomic replication elongation[57].

**Supplementary Figure 4.**

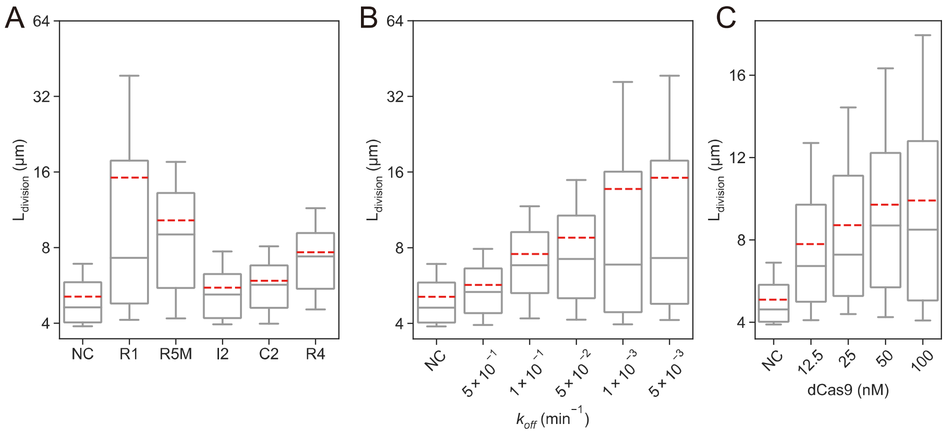

**Supplementary Figure 4. Simulated division lengths as the strength of *CRISPRori* was tuned by target site identities (A), dCas9 expression levels (B) and first order dCas9-sgRNA-DNA dissociation rates (C).** Unless otherwise indicated, the default parameters were: dCas9 concentration = 100 nM, $k_{\mathrm{off}}$ = 0.005 $\mathrm{min}^{-1}$. Target boxes are R5M and R1 for (B) and (C), respectively. For all groups, $N_{\mathrm{cell}}$ = 100.

**Supplementary Figure 5.**

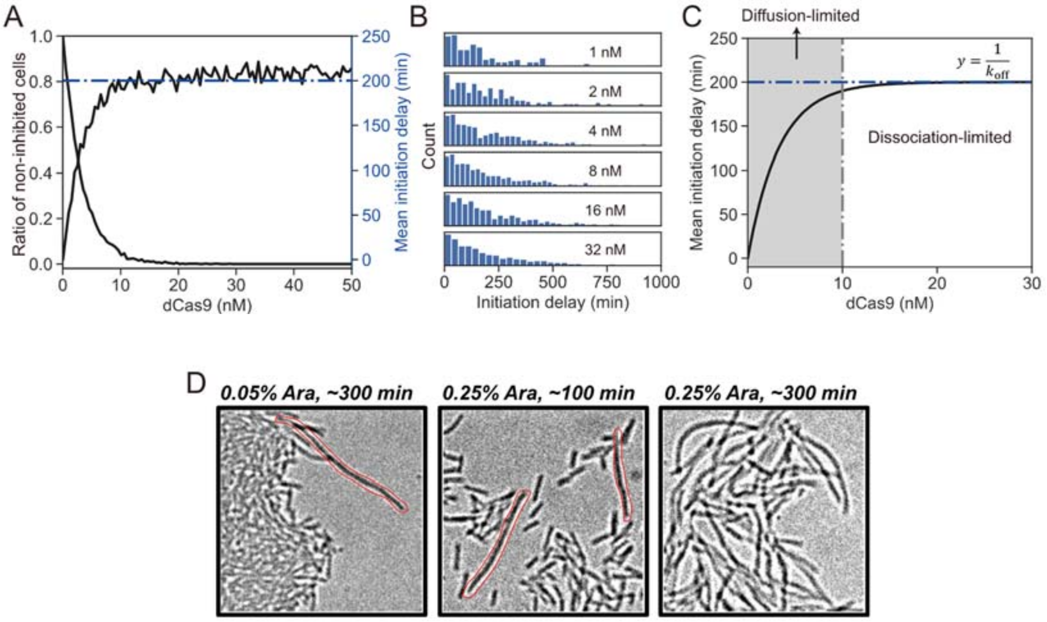

**Supplementary Figure 5. Phenotypic divergence of *CRISPRori* perturbed cells.** (A) The process of dCas9 association and dissociation are simulated by Gillespie algorithm. 1,000 simulations were performed for each concentration of dCas9. The output is the ratio of the cells where dCas9 failed to bind to *oriC*, and the mean delay of all repeats. (B) Distribution of initiation delay under variant dCas9 concentrations. Non-inhibited cells are not included here. The similarities among the distributions suggest that dCas9 dissociation is the only major factor that determines the inhibitory strength as long as dCas9 successfully binds to oriC. (C) Theoretical curve of mean delay, which converges to the reciprocal of dCas9's dissociation rate. (D) Cells under *CRISPRori* treatment with low inducer concentration (left) or in the early stage of induction (middle) shows higher morphological heterogeneity, compared to the late stage of high-concentration inducer condition (right).

**Supplementary Figure 6.**

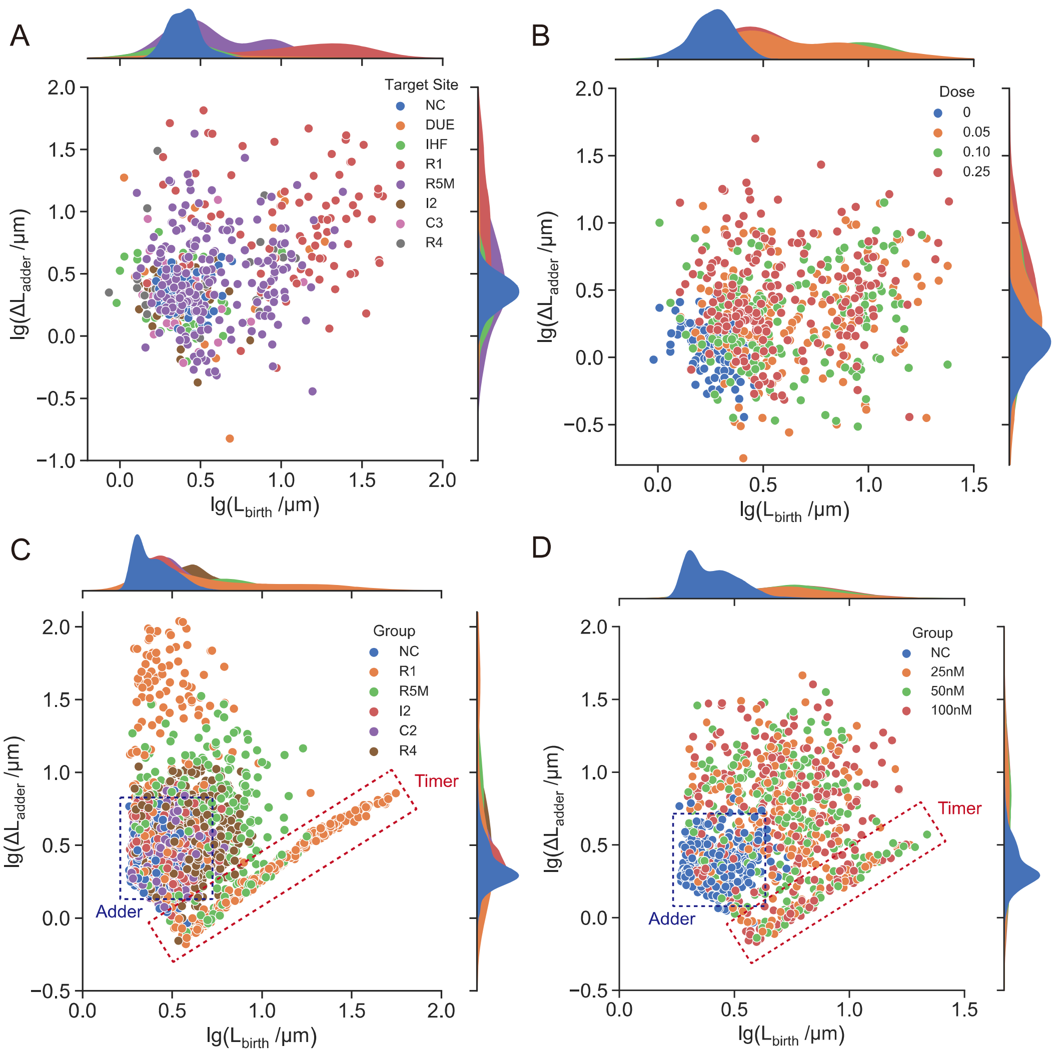

**Supplementary Figure 6. The breakdown of the division adder in *CRISPRori* perturbed cells.** (A) and (B) are experimental results, while (C) and (D) are corresponding simulated results.

**Supplementary Figure 7.**

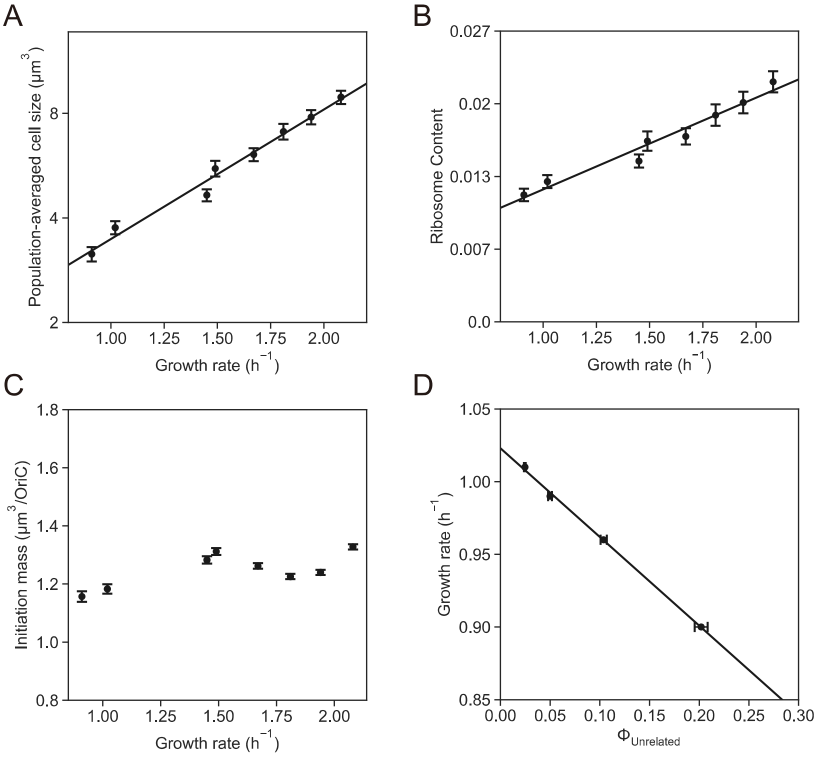

**Supplementary Figure 7. Model simulations of normally replicating cells conformed to several classic growth laws.** For (A), (B) and (C), The intracellular amino acid concentration is varied to generate different growth rates. (A) Schaechter-Maaloe-Kjeldgaard (SMK) growth law, states that population-averaged cellular mass scales exponentially with the growth rate. (B) Ribosome content increases linearly with the growth rate. (C) Donachie's constant-initiation-mass law, states that the cell mass per *oriC* at DNA replication initiation is a growth-rate-independent constant number. (D) The growth rate decreases linearly with ratios of unrelated proteins. The plasmid copy number is varied to generate different ratios of unrelated proteins. **All error bars are 95% C.I. Growth rates are calculated by linear fitting v to dv/dt.**

**Supplementary Figure 8.**

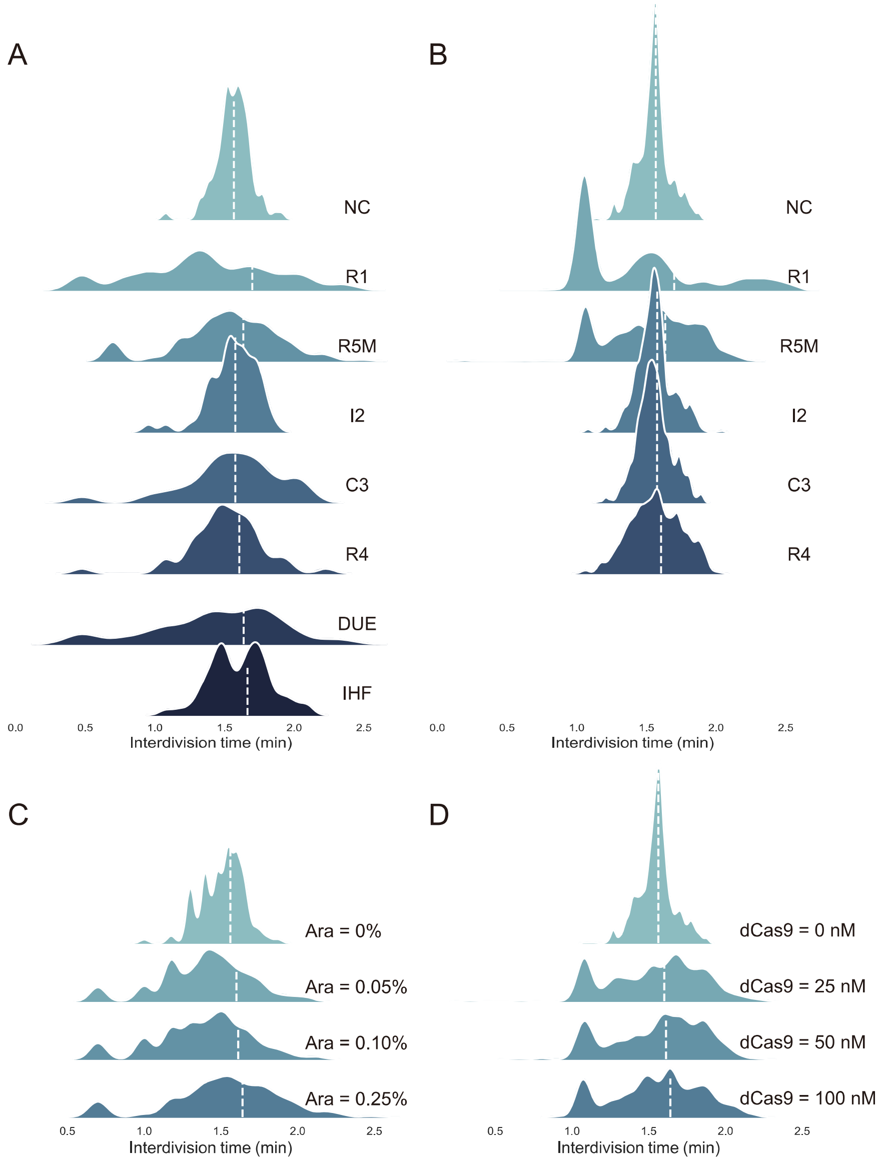

**Supplementary Figure 8. Interdivision time of *CRISPRori* perturbed cells in different groups.** (A) and (C) are experimental results, while (B) and (D) are corresponding simulated results.

**Supplementary Figure 9.**

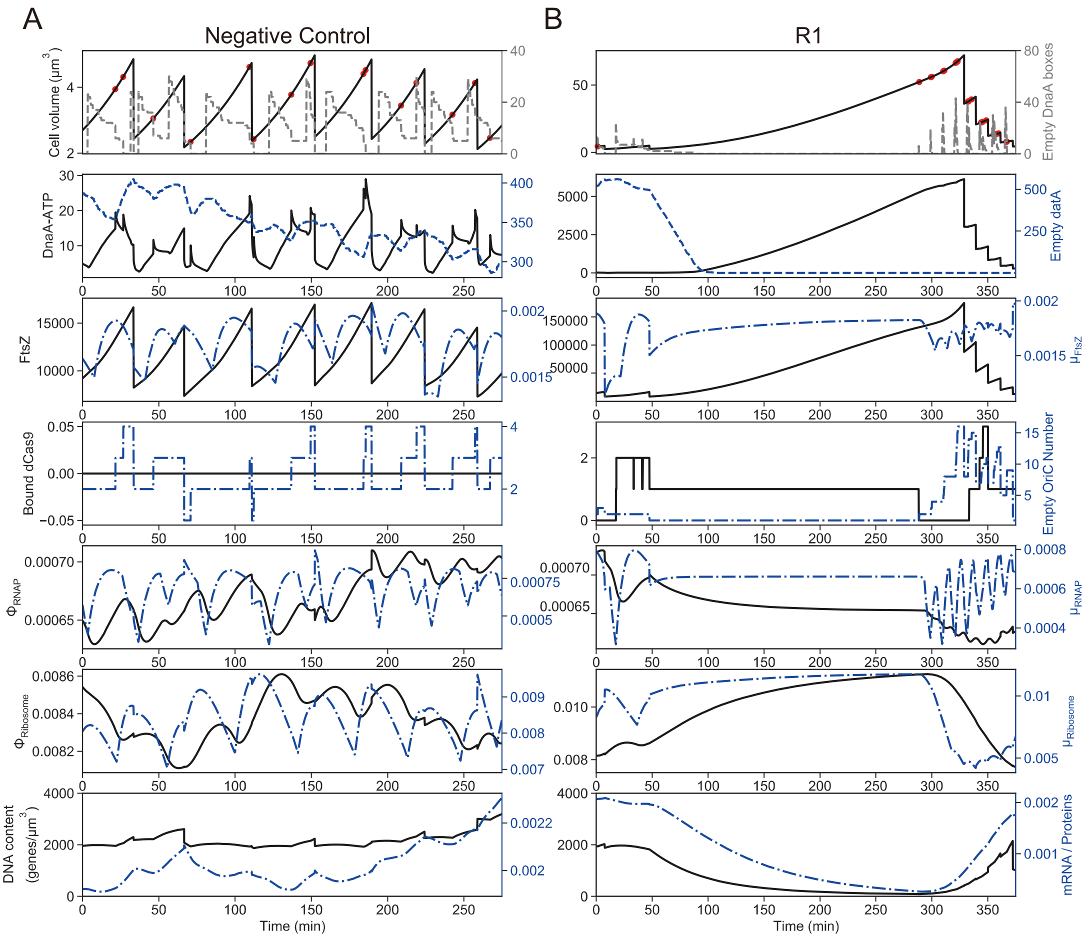

**Supplementary Figure 10. Simulated lineage dynamics of normally replicating cells and *CRISPRori* perturbed cells.** μ and Φ are the mRNA fraction and the proteomic fraction, respectively.

**Supplementary Figure 10.**

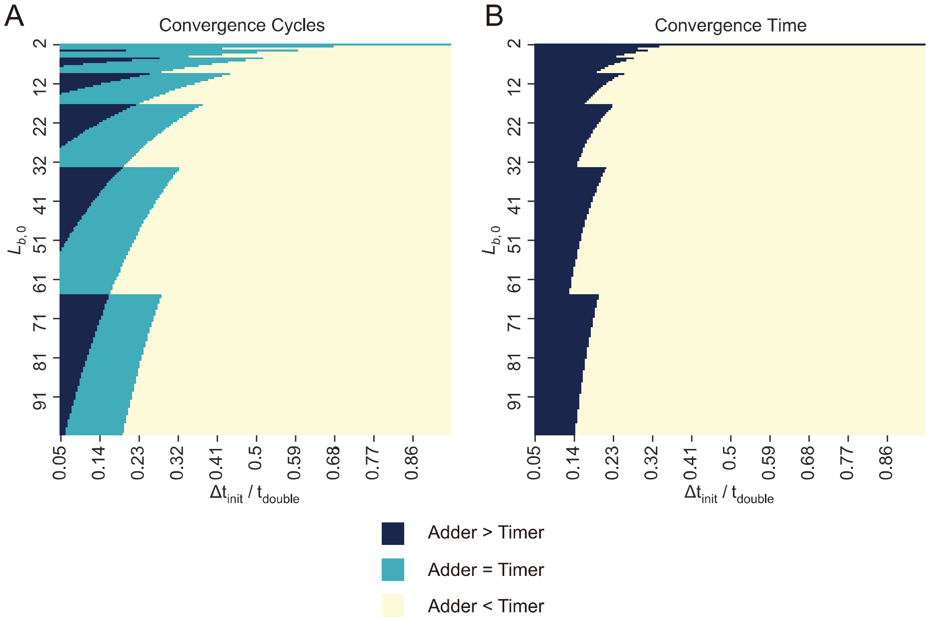

**Supplementary Figure 9. Comparison of number of division cycles (A) and time (B) for an elongated cell ($L_{birth} = n\Delta L_{adder}$, n > 2) to converge to normal range ($L_{birth} < 2\Delta L_{adder}$), between adder and timer.**

**Supplementary Figure 11.**

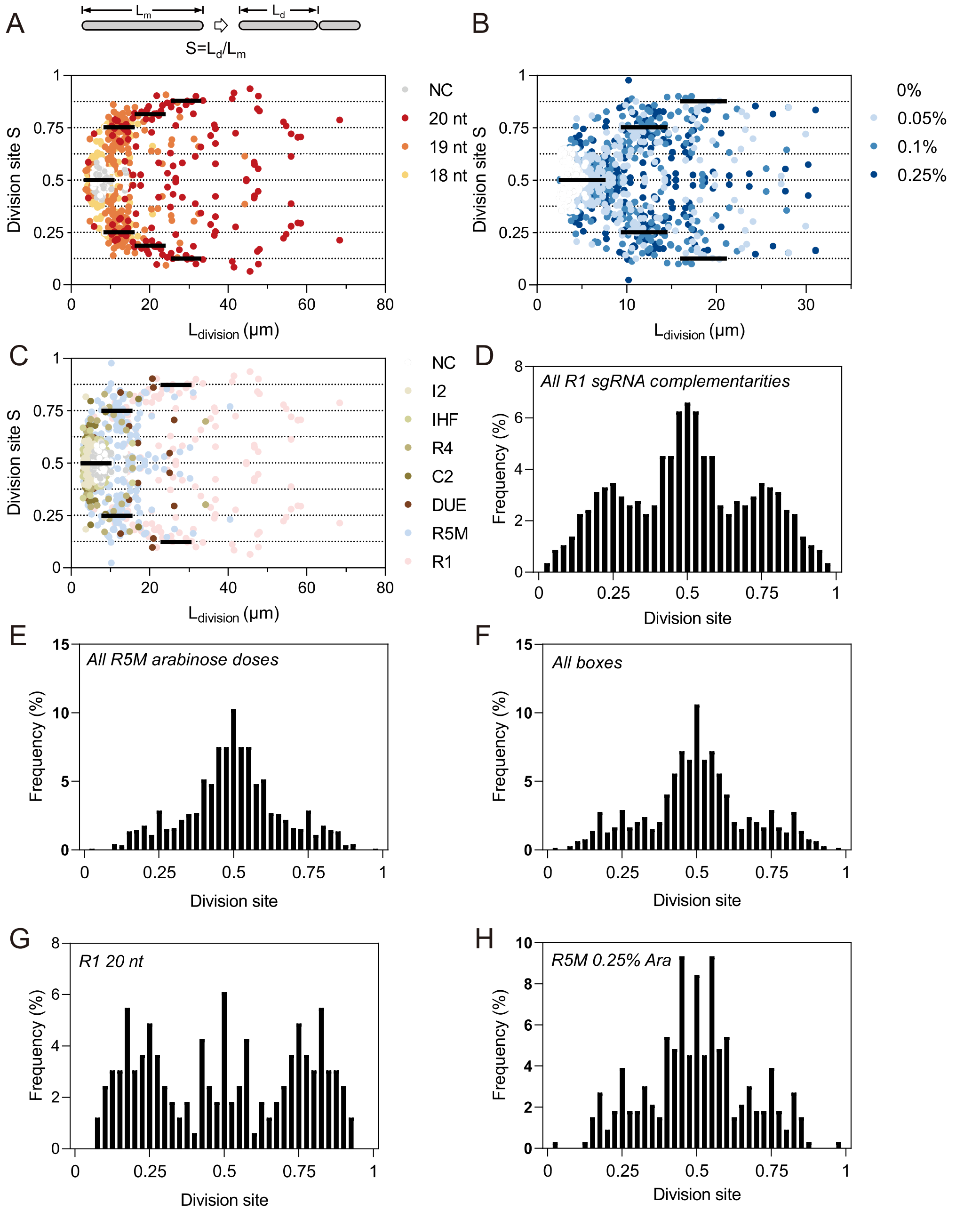

**Supplementary Figure 11. Division locations in *CRISPRori* perturbed cells.** Increased noise occasionally led to extremely small, nucleoid-free daughter cells.

**Supplementary Figure 12.**

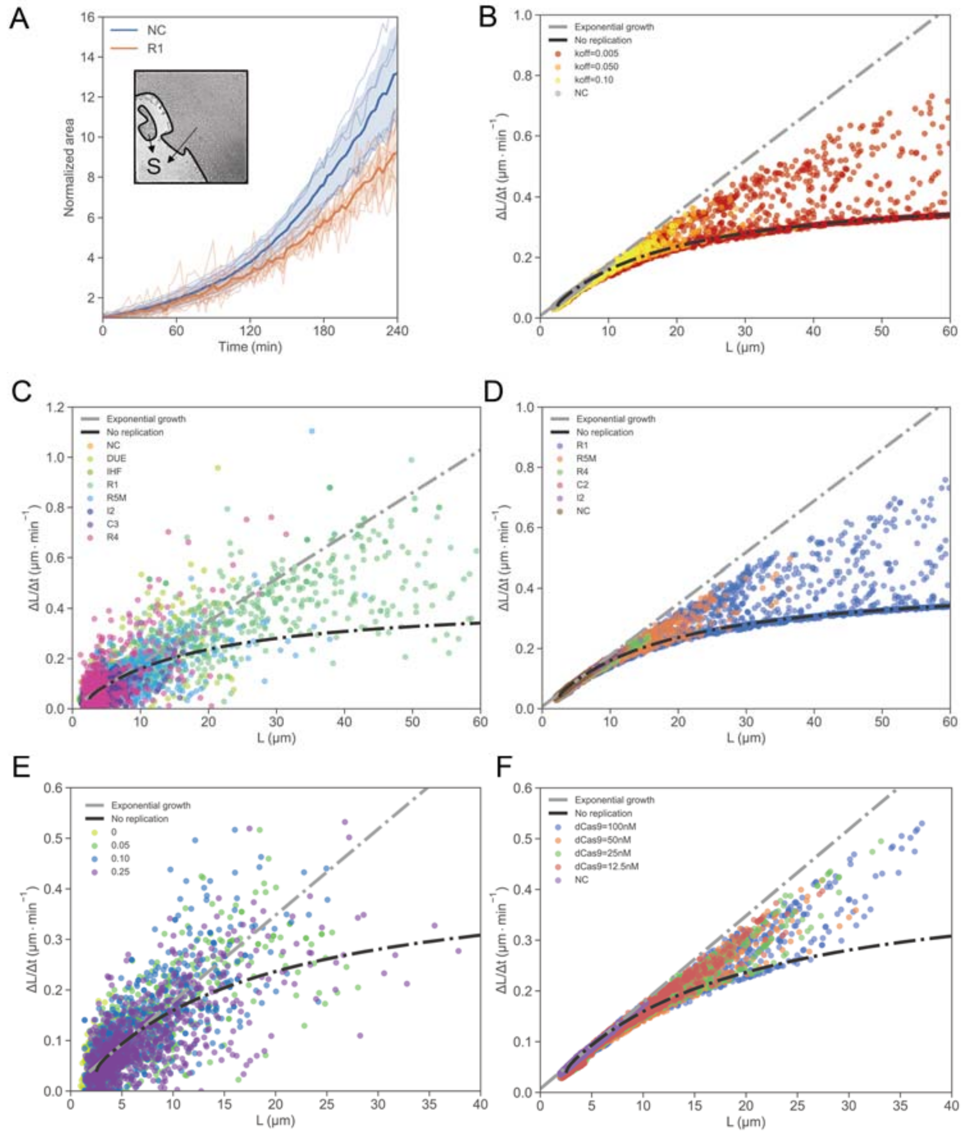

**Supplementary Figure 12. Transition from exponential to linear growth under *CRISPRori* perturbation.** (A) Bulk (population) growth rate change in *CRISPRori* perturbed cells (target = R1 box). Cell growth was characterized by the total area covered by cells in a microfluidic chamber (inset). The negative control group showed an exponential growth pattern. For the *CRISPRori* perturbed groups, linear growth was observed at >100 min. Shaded areas showed the 95% C.I. of all trajectories. $N = 7$ for both groups. (B)-(F) Single cell growth rates distributed differentially between exponential and linear growth modes for *CRISPRori* perturbed cells. (C) and (E) are experimental results, while (B), (D) and (F) are simulated results.

**Supplementary Figure 13.**

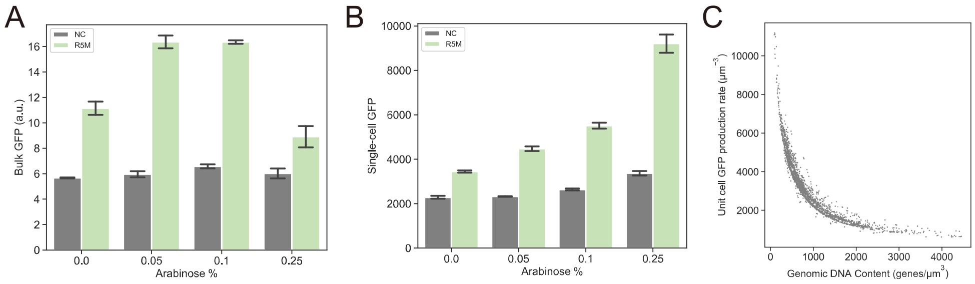

**Supplementary Figure 13. Enhanced expression of plasmid encoded GFP by genomic replication initiation perturbation.** (A) Batch-culture fluorescence as measured on plate reader. Total fluorescence peaked between 0.05% and 0.1% arabinose and subsequently decreased due to reduced total biomass. Error bars show the s.d. of three experimental replicates. (B) Single-cell fluorescence, as measured by flow cytometry, increased monotonously with perturbation strength. Error bars show the s.d. of three replicates, $N_{\mathrm{cell}}$ = 50000 for each. (C) Simulation of negative correlation between DNA content and unit-cell GFP production rate.

**Supplementary Figure 14.**

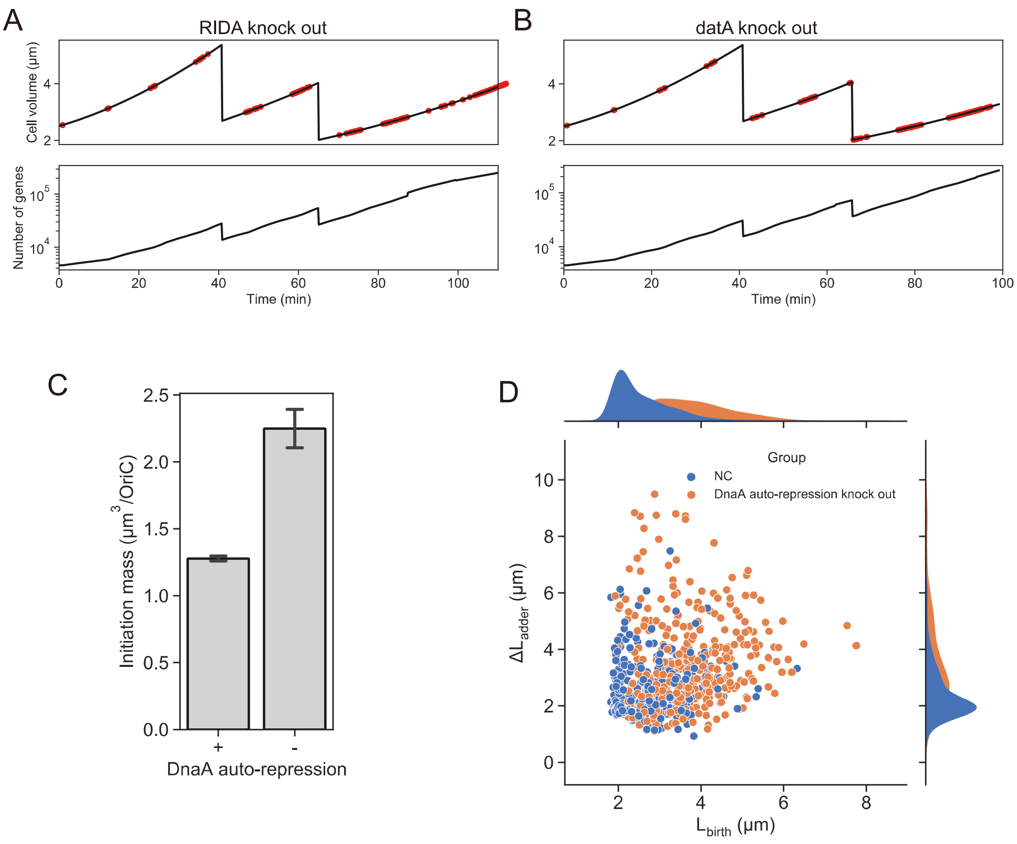

**Supplementary Figure 14. Simulated non-inhibition cells with several key negative regulations on DnaA-related replication initiation knocked out.** With a mechanism known as replicative inactivation of DnaA (RIDA) (A), and DnaA-titrated loci (datA) (B) removed from the cell, the genome undergoes severely excessive replication initiation. Eliminating the auto-repression of DnaA itself results in increase in initiation mass (C) and deviation from adder (D).
